# Common and divergent gene regulatory networks control injury-induced and developmental neurogenesis in zebrafish retina

**DOI:** 10.1101/2023.08.08.552451

**Authors:** Pin Lyu, Maria Iribarne, Dmitri Serjanov, Yijie Zhai, Thanh Hoang, Leah J. Campbell, Patrick Boyd, Isabella Palazzo, Mikiko Nagashima, Nicholas J. Silva, Peter F. HItchcock, Jiang Qian, David R. Hyde, Seth Blackshaw

## Abstract

Following acute retinal damage, zebrafish possess the ability to regenerate all neuronal subtypes. This regeneration requires Müller glia (MG) to reprogram and divide asymmetrically to produce a multipotent Müller glia-derived neuronal progenitor cell (MGPC). This raises three key questions. First, does loss of different retinal cell subtypes induce unique MG regeneration responses? Second, do MG reprogram to a developmental retinal progenitor cell state? And finally, to what extent does regeneration recapitulate retinal development? We examined these questions by performing single-nuclear and single-cell RNA-Seq and ATAC-Seq in both developing and regenerating retinas. While MG reprogram to a state similar to late-stage retinal progenitors in developing retinas, there are transcriptional differences between reprogrammed MG/MGPCs and late progenitors, as well as reprogrammed MG in outer and inner retinal damage models. Validation of candidate genes confirmed that loss of different subtypes induces differences in transcription factor gene expression and regeneration outcomes. This work identifies major differences between gene regulatory networks activated following the selective loss of different subtypes of retina neurons, as well as between retinal regeneration and development.

## INTRODUCTION

The zebrafish retina possesses a remarkable ability to regenerate neurons lost to acute damage by injury-induced reprogramming of Müller glia (MG) ^1^. A variety of damage models have been used to study zebrafish retinal regeneration that show varying levels of specificity to neuronal subtypes lost. These include rod and cone photoreceptor ablation using bright light ^2–4^, excitotoxic and ouabain-mediated destruction of retinal ganglion and amacrine cells ^5–7^, needle poke destroying all neuronal layers ^8^, heat damage ^9^, and the use of cell-specific nitroreductase transgenes to achieve cell type-specific ablation ^5, 10–12;^. Regardless of the damage model utilized, reprogrammed MG divide asymmetrically to produce a Müller glia-derived neuronal progenitor cell (MGPC), which continues to proliferate ^2, 3, 13^. These MGPCs regenerate specific cell types that are lost to injury, and these MGPC-derived neural precursors in turn migrate to the appropriate retinal layers ^5, 10, 14, 15^. In every retinal injury model, MG rapidly upregulate genes specific to neural progenitors such as *ascl1a* and *lin28a* and begin proliferating within 36 hours following injury ^16–19^. This results in the efficient regeneration of all retinal neuronal subtypes within several weeks following initial injury^2, 3, 15^.

Two major models have been proposed to explain regeneration of different neuronal subtypes in the retina. One model proposes that MGPCs respond to the loss of specific neuronal subtypes and are biased to primarily commit and differentiate into those lost neurons in a damage-dependent model ^5, 14, 20, 21^. The alternative model suggests that MG reprogram to a state identical to retinal progenitors found in development and the MGPCs then progress through regeneration in a process mimicking retinal development ^1, 15^, with MGPC-derived neuronal subtypes that are not lost to injury failing to integrate efficiently into retinal circuitry and thereafter being lost. Several groups have examined how similar regeneration is to retinal development utilizing the temporal expression of developmental genes during regeneration ^14, 15, 22^. However, these results are limited based on the number of genes and time points examined. In addition, a systematic comparison of the fate of MGPC-derived neurons produced in different injury models has yet to be performed, which is critical for understanding the potential for common features across retinal regeneration.

In this study, we systematically examined the similarities and differences between two different cell-selective models of acute damage: bright light-induced destruction of rod and cone photoreceptors and NMDA-mediated excitotoxicity that results in loss of amacrine and ganglion cells. Using long-term EdU-based fate mapping, we found that MGPC-derived neurons generated following these two different injuries are not strictly cell type-specific, although they do show a consistent bias towards neurons that are preferentially lost following injury. We found that considerable numbers of inner retinal neurons were generated following light damage, and likewise many photoreceptors following NMDA injury, and these newly generated neurons are not later eliminated by selective cell death.

To investigate the molecular mechanisms controlling injury-induced neurogenesis, we performed both single-cell RNA-Seq (scRNA-Seq), single-nuclei RNA-Seq (snRNA-Seq), and ATAC-Seq multiomic analysis to comprehensively profile injury-induced changes in gene expression and chromatin accessibility seen in both injury models. We sought to determine how closely these MGPCs resembled retinal progenitor cells (RPCs) in developing retinas. We observed that MGPCs produced in both retina damage models exhibit similarities and differences in gene expression and gene regulatory networks that could account for biases in neurogenesis, and we also identified the secreted metalloprotease *mmp9* as a selective inhibitor of amacrine and ganglion cell formation. Though we found that activated MG reprogrammed to a state similar to late-stage RPCs in developing retinas, there were distinct differences that existed between these cell types, with development and regeneration possessing several unique gene regulatory networks. We further identified the transcription factor (TF) *foxj1a* as essential for injury-induced neurogenesis. These data demonstrate that retinal regeneration is similar to, but does not precisely recapitulate, retinal development.

## RESULTS

### Light damage and NMDA-induced excitotoxicity result in regeneration of overlapping neuronal cell subtypes

It was previously shown that constant intense light results in the loss of rod and cone photoreceptors ^2^, while NMDA damage results in the loss of amacrine and ganglion cells ^5^. Both damage models induce MG reprogramming, MGPC production and proliferation, and regeneration of retinal neurons. However, a careful analysis of the extent of regenerated neuronal subtypes in these two injury models has not yet been performed. We labeled proliferating MGPCs using EdU incorporation from 60 to 108 hours following both NMDA damage and light damage (LD), examined the neuronal cell subtypes that incorporated the label following NMDA damage, and compared these to uninjured controls (Fig. 1A,B). All cell counts are displayed in Table ST1. While we detected extensive EdU incorporation in amacrine and ganglion cells at both 7 (7DR) and 14 (14DR) days following NMDA injury, we also observed extensive EdU incorporation into cells throughout both the inner nuclear layer (INL) and the photoreceptor-dominated outer nuclear layer (ONL) at these time points (Fig. 1B,D). No change in either the total number of EdU-positive cells was detected between 7 days of recovery (7DR) and 14DR (Fig. 1C), or in the number of EdU-labeled ONL cells between 7DR and 14DR (Fig. S1A,B). Most EdU-positive neurons were found in the INL and ganglion cell layer (GCL) at both timepoints, with 62.5% of EdU-positive cells at 7DR and 60% of EdU-positive cells at 14DR (Fig. 1E). The remaining EdU-labeled cells were present in the ONL at both time points (Fig. 1E, 37.5% and 40%). Uninjured controls possessed low numbers of EdU-labeled cells in the ONL only, corresponding to adult-born rod photoreceptors as previously reported ^5, 23, 24^ (Fig. 1B-D).

**Figure 1:**
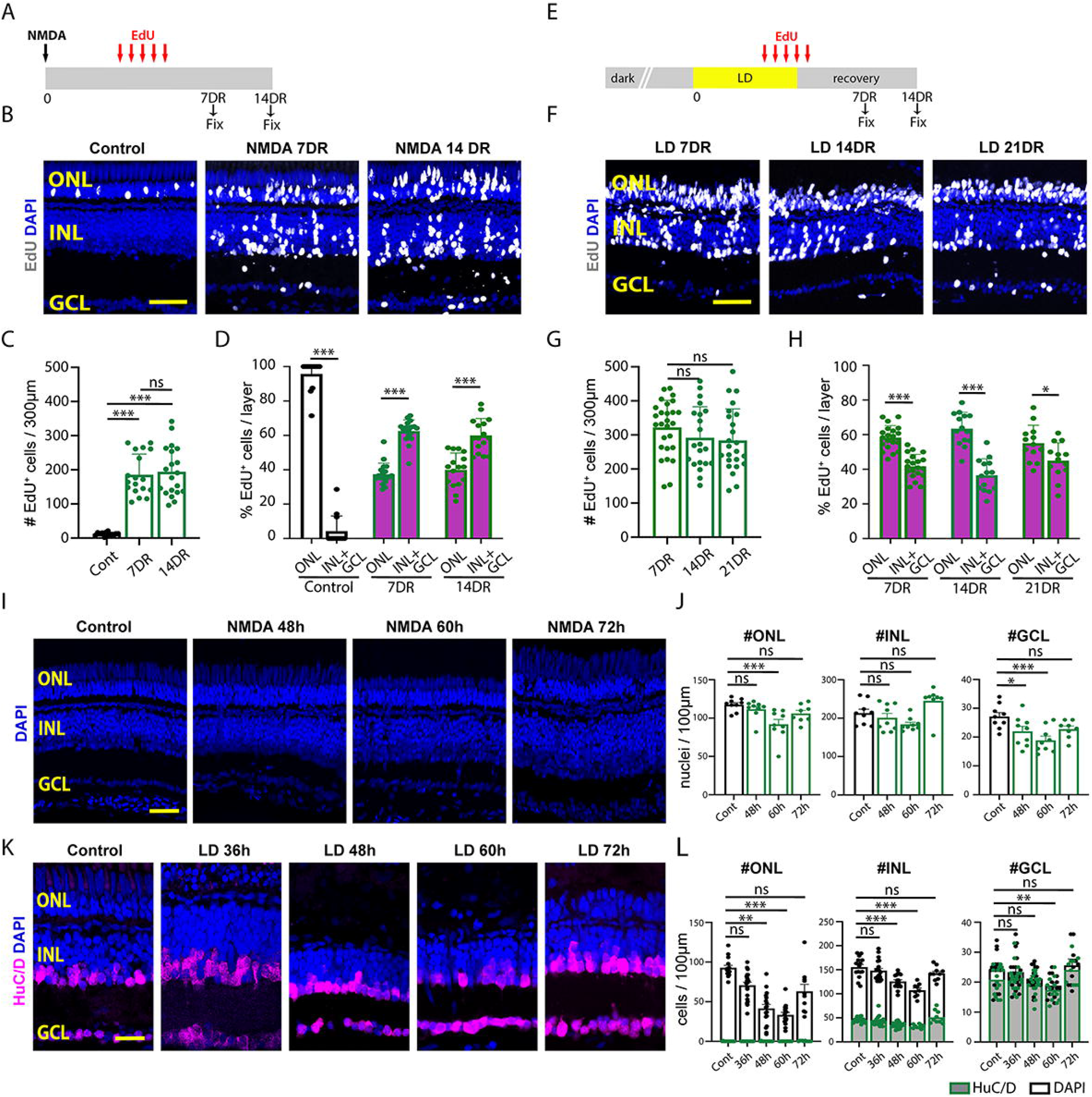
Comparison of NMDA-induced and light-induced retinal damage. (A) Schematic of NMDA-induced damage experiment. (B) EdU-labeling in control retinas and following NMDA damage. (C) Quantification of the number of EdU-labeled cells in all three retinal layers. (D) The percentage of the EdU-labeled cells in the Outer Nuclear Layer (ONL) versus combined in the Inner Nuclear Layer (INL) and Ganglion Cell Layer (GCL). (E) Schematic of light-induced damage experiment. (F) EdU-labeling following light damage. (G) Quantification of the number of EdU-labeled cells in all three retinal layers. (H) The percentage of the EdU-labeled cells in the ONL versus combined in the INL+GCL. (I) DAPI staining of undamaged retinas and 48, 60, and 72 hours after injecting NMDA. (J) Quantification of the number of DAPI-labeled nuclei in the ONL. (K) Quantification of the number of DAPI-labeled nuclei in the INL. (L) Quantification of the number of DAPI-labeled nuclei in the GCL. (M) DAPI and HuC/D staining of undamaged retinas and 36, 48, 60, and 72 hours after starting constant light treatment. (N) Quantification of the number of DAPI-or HuC/D-labeled nuclei in the ONL. (O) Quantification of the number of DAPI-or HuC/D-labeled nuclei in the INL. (P) Quantification of the number of DAPI-or HuC/D-labeled nuclei in the GCL. Scale bars in B, F, and I are 20μm and in M is 14μm.

Using cell-specific immunohistochemical markers, we observe a slight increase in the number of both EdU/HuC/D-positive amacrine and ganglion cells (Fig. S1A,C) and EdU/Zpr1-positive double cone photoreceptors (Fig. S1F,H) from 7DR to 14DR. In contrast, there is a significant decrease in the number of EdU/PKCα-positive bipolar cells (Fig. S1A,D) and a slight decrease in the number of EdU/4c12-positive rod photoreceptors (Fig. S1E,G) between 7DR and 14DR, indicating possible removal of excess immature neurons. There was no significant difference in the number of EdU/GFAP-positive MG (Fig. S1I,K) and EdU-labeled microglia/macrophages (Fig. S1J,L) from 7DR to 14DR. These results indicate that MGPCs induced by both LD and NMDA generate primarily photoreceptors, amacrine and ganglion cells. Furthermore, with the exception of a small number of bipolar cells and rod photoreceptors, no selective loss of newly generated neurons was detected between 7 and 14 days following NMDA damage, and a large percentage of regenerated neurons were photoreceptors, which were previously thought not to be significantly damaged by NMDA.

We next examined the regenerated retina at 7, 14, and 21 days recovery following light damage, and also observed extensive EdU incorporation in all cell layers (Fig. 1F, S2A). As with the NMDA-injured retina, no change in the total number of EdU-positive cells was detected between 7DR and 21DR (Fig. 1G), or in the total number of EdU-labeled INL cells between (Fig. S2A,B). In contrast to the NMDA-damaged retina, approximately 60% of all EdU-positive cells were located in the ONL at all time points (58% at 7DR, 63% at 14DR, and 55% at 21DR (Fig. 1F, H; S2), with approximately 40% of the EdU-positive nuclei located in the INL and GCL. EdU-positive cells in the ONL were extensively co-labeled with the rod photoreceptor marker 4c12 (Fig. S2A,C), while EdU-positive cells in the INL were predominantly HuC/D-positive (Fig. 1L, S2E,F). As with NMDA injury, few EdU-positive cells colabeled with either the MG marker GFAP (Fig. S2A,D) or the bipolar cell marker PKCα (Fig. S2E,G), while essentially no EdU-positive horizontal cells were detected (data not shown). Again, the total number of EdU-labeled cells remained statistically the same for all cell type-specific markers tested from 7DR through 21DR (Fig. 1G, Fig. S2), demonstrating that there was no selective loss of any particular neuronal subtype in this period.

To examine the reason for this widespread EdU incorporation in both damage models, we carefully examined the loss of retinal neurons in both damage models. During NMDA damage, we quantified the number of DAPI-stained nuclei in all three nuclear layers in undamaged (control) and NMDA-damaged retinas at 48, 60, and 72 hours following NMDA injection (Fig. 1I,J). As expected, there is a significant reduction in the number of GCL nuclei relative to control, but not a significant reduction in the number of INL nuclei (Fig. 1J). However, there is also an unexpected and significant reduction in the number of ONL nuclei at 60 hours following NMDA injection relative to the control (Fig. 1J). Likewise, in light-damaged retinas, a significant loss of ONL nuclei was detected by 36 hours, with lowest numbers seen at 60 hours, and a recovery to baseline levels observed by 72 hours post-injury (Fig. 1K,L). However, we also observed significant reductions in the number of nuclei in both the INL and GCL by 48 and 60 hours following light damage, respectively (Fig. 1K,L). Thus, both NMDA and constant light result in the loss of retinal cell types (photoreceptors and inner retinal neurons, respectively) that were unexpected and could account for the regenerated neurons in those layers.

### ScRNA-Seq and snRNA/ATAC-Seq multiomic analysis of NMDA- and light-damaged retina

To comprehensively characterize changes in gene expression and regulation that occur following both NMDA and light-mediated injury, we performed both scRNA-Seq and combined multiomic snRNA/ATAC-Seq analysis of whole retina at multiple timepoints following injury. We analyzed uninjured control samples along with retinas harvested at 36hr, 54hr, 72hr, 96hr, 7days, and 14 days post-injury (Table ST2). In total, we profiled 99,555 cells using scRNA-Seq and 137,490 cells using snRNA/ATAC-Seq multiomic analysis. UMAP analysis of integrated datasets from both scRNA-Seq and snRNA/ATAC-Seq multiomic analysis readily identified all major adult retinal cells in both uninjured control and each condition examined (Fig. 2A,B; S3A,B). In addition, specific cell subtypes were detected in both injury models that were either absent or present at much lower levels in uninjured controls. These include activated MG, MGPCs and immature precursors of rod and cone photoreceptors, amacrine and ganglion cells (Fig. 2C). More rod and cone precursors were consistently detected following light damage, while more RGC precursors were detected following NMDA treatment (Fig. 2D, S3C). Relatively few bipolar and no horizontal cell precursors were detected, matching the observed EdU data in Fig. 1. Well-characterized cell type-specific markers showed both expected patterns of gene expression and chromatin accessibility as determined by both scRNA-Seq and snRNA/ATAC-Seq multiomic analysis (Fig. S3D Table ST2).

**Figure 2:**
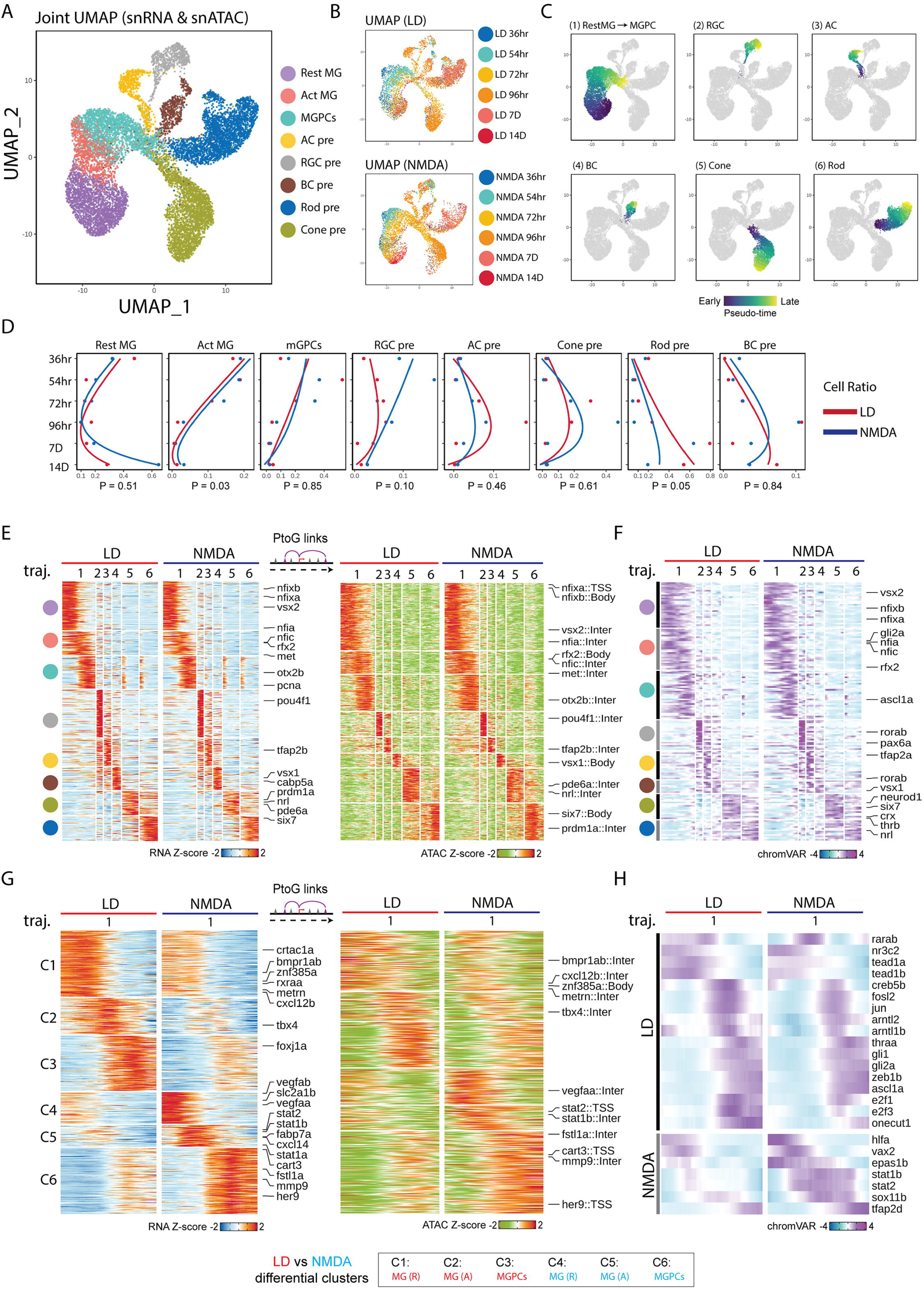
Shared and differential patterns of gene expression and chromatin accessibility data observed in MG-derived cells following LD and NMDA treatment. (A,B) Combined UMAP projection of MG and progenitor neuron cells profiled using multiomic sequencing. Each point (cell) is colored by cell type (A) and time points (B). (C) UMAPs showing trajectories constructed from mutiomics datasets of combined LD and NMDA datasets. Color indicates pseudotime state. (D) Line graphs showing the fraction of cells (x axis) at each time point (y axis) of each cell type. Lines are colored by treatment. (E) Heatmap shows the consensus marker genes and their related marker peaks (TSS and enhancer) between LD and NMDA treatment for each cell type. (F) Heatmap shows the consensus motifs between LD and NMDA treatment for each cell type. (G) Heatmap shows the differential genes and their related differential peaks (TSS and enhancer) between LD and NMDA treatment for MG(R), MG(A) and MGPCs. (H) Heatmap shows the differential motifs between LD and NMDA treatment for MG(R), MG(A), and MGPCs.

Analyzing the integrated snRNA/ATAC-Seq dataset (Fig. 2A), we observe dynamic changes in cell composition across each timepoint for both injury models (Fig. 2B-D). We notice a progressive reduction in the relative fraction of activated MG and MGPCs from 36hr onward, while the majority of MG return to a resting state by 14 days following injury (Fig. 2D). A modest but significant increase in the relative fraction of activated MG was observed following NMDA injury relative to light damage (Fig. 2D).

The fraction of RGC precursors likewise decreases from 36 hours following NMDA injury but declines more slowly following light damage, while the fractions of amacrine and cone photoreceptor precursors peak at 96 hours following both NMDA and light damage (Fig. 2D). Finally, the fraction of rod photoreceptor and bipolar precursors steadily increases with time for both injury models (Fig. 2D). This broadly corresponds to the observed order of neurogenesis in the developing zebrafish retina, where ganglion cells are born first and bipolar cells and rod photoreceptors last ^25, 26^, and also aligns with results obtained from a recent scRNA-Seq analysis of MGPC and MGPC-derived neurons obtained from light-damaged retina ^27^.

To identify genes that control both injury-induced MG reprogramming and specification and differentiation of MGPC-derived neurons, we performed pseudotime analysis of the integrated snRNA/ATAC-Seq dataset from both NMDA and light damage samples (Fig. 2C). We identified six distinct trajectories, corresponding respectively to the transition from resting MG to activated MG to MGPCs; from MGPCs to RGCs, amacrine cells, bipolar cells, cone photoreceptors, and rod photoreceptors (Fig. 2C). Dynamic patterns of differential accessibility, which were detected using scATAC-Seq analysis, were observed for both promoter and putative distal regulatory elements, with both enhancer and silencer elements detected (Fig. 2E). A total of 4,891 dynamically expressed genes and 13,988 differentially accessible DNA elements were observed in both NMDA and light-damaged samples (Table ST3).

Within the first trajectory, three separate sequential elements were detected, corresponding to genes selectively expressed in resting MG, activated MG, and MGPCs respectively (Fig. 2E, Table ST3). Resting MG expressed TFs known to mediate glial quiescence such as *nfia*, activated MG expressed TFs such as *vsx2* and *foxj1a*, while MGPCs expressed *atoh7* and cell cycle regulators such as *mki67*. Each trajectory included multiple TFs previously shown to be essential for specification of the respective cell type *pou4f1* in RGCs (2), *tfap2c/d* in amacrine cells (3), *vsx1* in bipolar cells (4), *nrl* in rods (5), and *six7* in cone photoreceptors (6). Gene Ontology analysis revealed distinctive features of genes enriched in each trajectory, with genes specific to regulation of neurogenesis enriched in activated MG, cell cycle and ribosomal genes enriched in MGPCs, and axonogenesis enriched in immature RGCs (Fig. 2F). We also analyzed the scATAC-Seq data for differentially accessible TF binding motif sequences, and identified 185 differential motifs (Fig. ST3). These likewise include target sites for most differentially expressed TFs both NMDA and light damage, including *foxj1a*, *tfap2b* and *nrl* (Fig. 2G,H).

### NMDA and light damage differentially regulate gene expression in MG and MGPCs

These data show that while NMDA-mediated excitotoxicity and light damage induce formation of MGPCs through broadly overlapping mechanism, there are nonetheless important quantitative differences seen in both gene expression and regulation in these two injury models. To identify differences in gene expression and regulation in MG following NMDA and light damage, we directly compared differences in mRNA expression, and both chromatin and motif accessibility in the two injury models. While we identified 4,891 genes that showed similar patterns of expression in both light damage and following NMDA treatment, we also identified 1,791 genes showing higher expression in light damage and 319 following NMDA (Fig. 2G) in MGs and MGPCs.

These include several genes that have been previously investigated in the context of developmental and/or injury-induced neurogenesis ^28–33^. Specifically, we observe *bmpr1ab*, *metrn*, *crtac1a*, and *cxcl12b* selectively and rapidly upregulated in activated MG of light-damaged retina, while *vegfaa/b*, *stat1a/b, stat2*, *cxcl14*, *her9*, *fstl1a*, and *mmp9* are rapidly upregulated following NMDA treatment (Fig. 2A,B; Table ST3). We also observe more pronounced upregulation of *foxj1a*, a TF which directs formation of multiciliated epithelial cells ^34, 35^, and *tbx4* following light damage. Both Gene Ontology and KEGG pathway analysis identified functional differences in differentially expressed genes between the two injury conditions, with genes associated with cell morphogenesis and adhesion enriched following light damage and genes controlling stress and immune response, as well as amino acid biosynthesis, enriched following NMDA treatment (Fig. S4A,B).

We likewise identified 1,052 chromatin regions that showed similar patterns of differential accessibility in both injury models, 811 chromatin regions that showed higher expression following light damage and 240 that showed higher expression following NMDA treatment in the snATAC-Seq dataset (Fig. 2G, Table ST3). Analysis of differentially accessible motifs revealed that 184 motifs showed shared patterns of differential accessibility in both injury models, but that 17 motifs were selectively enriched following light damage in MG and/or MGPCs, while 12 were enriched following NMDA treatment, with most showing relatively modest differences. Light damage induced increased accessibility at tead1, nr2e1, rxraa, and arntl1b motifs, while NMDA induced increased accessibility at stat1a/b, stat2, rfx2, and vax2 motifs (Fig. 2H).

### Mmp9 selectively suppresses regeneration of amacrine and ganglion cells

It has been previously shown that Mmp9 protein acts to repress MGPC proliferation in the light-damaged retina, with the light-damaged *mmp9* mutant exhibiting an increased number of MGPCs relative to light-damaged controls, and also exhibiting a loss of regenerated cones relative to control retinas ^33^. However, we observed that *mmp9* expression was elevated in activated MG and MGPCs in the NMDA-damaged relative to the light-damaged retina (Fig. 3A), and that NMDA treatment led to increased accessibility at putative cis-regulatory elements regulating *mmp9* transcription (Fig. 3B). This suggested that mmp9 might preferentially modulate the generation of inner retinal neurons, and we therefore examined the consequence of loss of *mmp9* expression in the regenerated retina in the two damage models, comparing the total number of EdU-labeled cells in undamaged control, NMDA-damaged, and light-damaged retinas at 7DR (Fig. 3C-G). We observed a significant increase in the total number of EdU-labeled neurons in the *mmp9* mutant relative to controls following light damage (Fig. 3D), as previously reported ^33^, which reflected an increase in the number of EdU-positive cells in both the ONL and INL (Fig. S5C,D). However, there was no significant increase in the total number of EdU-labeled neurons in the *mmp9* mutant following NMDA damage relative to controls (Figs. 3D, S5A,B). We also observed a significantly greater number of EdU-positive cells in both the INL and GCL in undamaged retinas, when the vast majority of newborn neurons are rod photoreceptors ^26^, as well as following light damage (Fig. 3E). However, this was not observed in NMDA damaged retinas. Following both light damage and NMDA treatment, moreover, we observe a significant increase in both the relative ratio of EdU-positive cells in the INL and GCL relative to the ONL (Fig. 3E), as well as an absolute increase in the number of HuC/D and EdU-positive amacrine and ganglion cells (Fig. 3F,G). All cell counts are listed in Table ST1.

**Figure 3:**
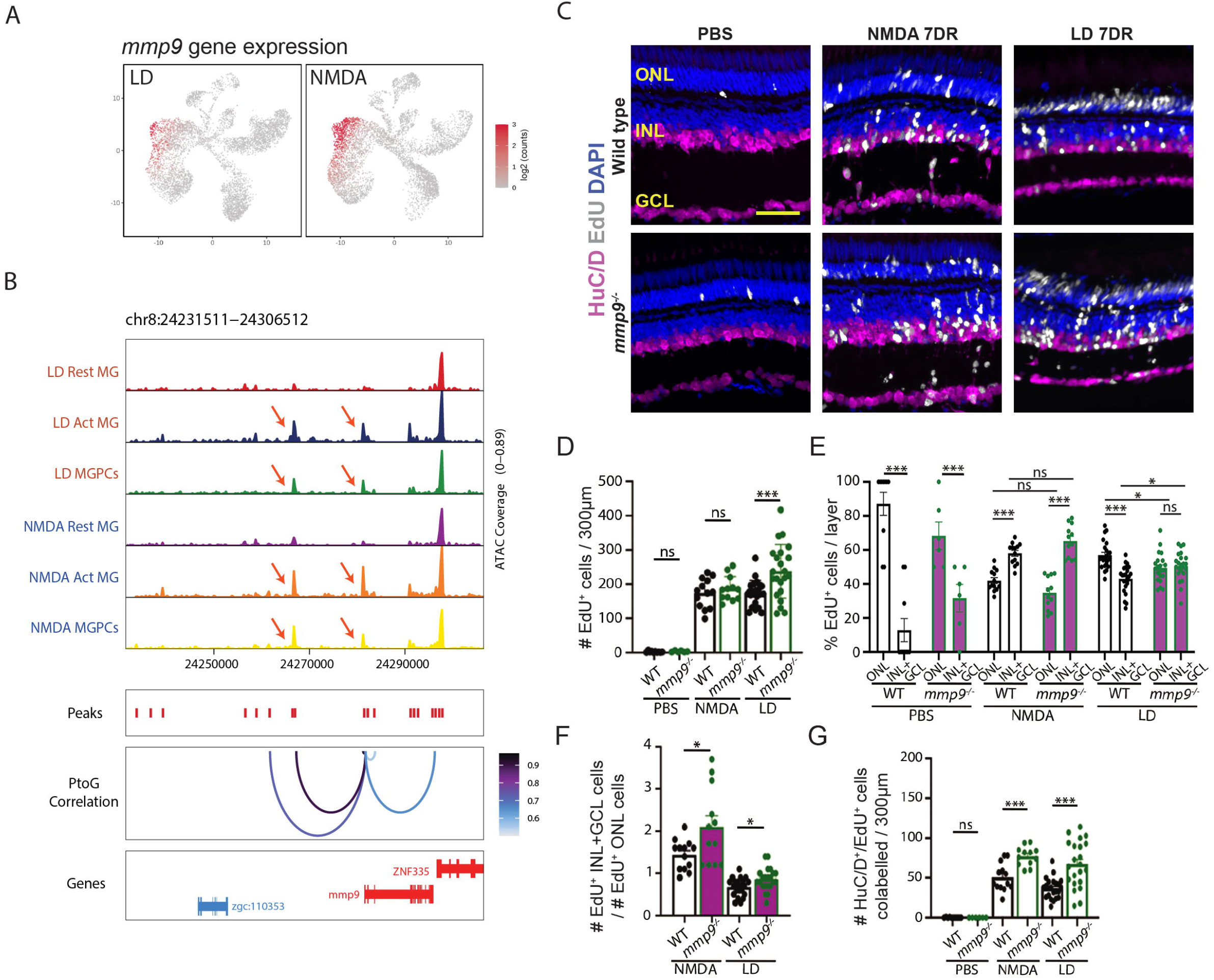
Mmp9 selectively inhibits generation of inner retinal neurons from MGPCs. (A) Gene expression pattern of mmp9 between LD and NMDA model. (B) Changing of chromatin accessibility around mmp9 loci between LD and NMDA models. (C) Schematic of NMDA-damaged experiments in wild-type and *mmp9* mutant fish. Retinas were PBS-injected (undamaged) or damaged for 96 hours and then isolated after 7 days of recovery. (D) Schematic of light-damaged experiments in wild-type and *mmp9* mutant fish. Fish were placed in constant darkness for 2 weeks and then exposed to constant intense light for 96 hours and then placed in standard light conditions and sacrificed 7 days later. (E) Retinal sections were immunostained with HuC/D and EdU and then counterstained with DAPI. (F) Quantification of the number of EdU-labeled cells in all three retinal layers in wild-type and *mmp9* mutants following either PBS injection, NMDA damage, or light damage. (G) The percentage of EdU-positive cells in the ONL versus the INL+GCL is plotted for wild-type and *mmp9* mutant retinas after PBS injection, NMDA damage, and light damage. (H) The ratio of EdU-positive ONL cells to EdU-positive INL+GCL cells is plotted for wild-type and *mmp9* mutant fish following either NMDA damage or light damage. (I) Quantification of the number of cells colabelled for EdU and HuC/D in PBS-injected, NMDA-injected, and light-damaged retinas. Scale bar in E is 20μm.

The relative increase in formation of generation of inner retinal neurons is already evident at 3DR in *mmp9* mutants following light damage, though not following NMDA treatment (Fig. S5E-J), possibly indicating the existence of additional factors inhibiting inner retinal neurons that initially compensate for loss of function of *mmp9*. The increased number of regenerated HuC/D-positive amacrine and ganglion cells in *mmp9* mutants relative to controls suggests that Mmp9 actively represses specification and/or differentiation of MGPC-derived amacrine and ganglion cells, in addition to its known role in repressing MGPC proliferation. To our knowledge, the *mmp9* mutant represents the first mutant that alters the commitment of MGPCs from one neuronal subtype (photoreceptors) to another subtype (inner retinal neurons).

### Identification of gene regulatory networks controlling differential injury response in NMDA and light-damaged MG

Having observed extensive differences in both gene expression and chromatin accessibility in activated MG and MGPCs following NMDA and light damage, we hypothesized that injury-induced gene regulatory networks are differentially organized in these two injury models (Fig. 4A). To test this, we used gene expression and chromatin accessibility data extracted from multiomic data to identify key transcriptional activators and repressors, finding that ∼60% of identified TF-Motif pairs act as transcription activators in both injury models (Fig. 4B). We next separately reconstructed injury-regulated transcriptional regulatory networks in MG and MGPCs following both NMDA treatment and light damage, identifying both TFs and target genes that directly target enhancer and/or promoter elements to activate or repress individual genes (Fig. 4C). We identified far more candidate activating relationships than repressive regulatory relationships in both injury models, with ∼85% of these targeting enhancers and ∼15% targeting promoters (Fig. 4C). We observe that 19% of all TF-peak-gene and 29% of all TF-gene regulatory relationships are common in the two injury models (Fig. 4D).

**Figure 4:**
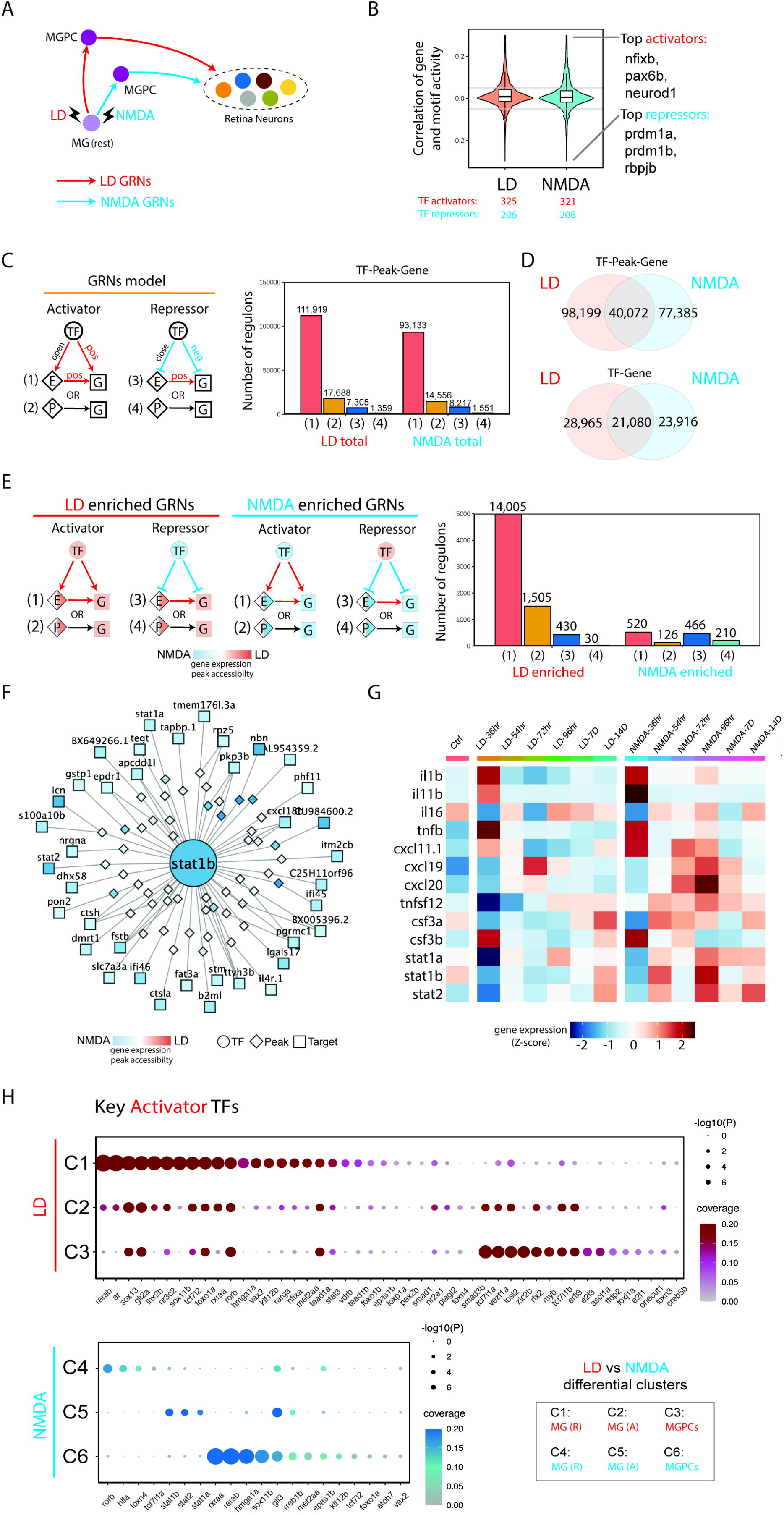
Transcription factors controlling differential gene expression in MG and MGPC following LD and NMDA treatment. (A) Schematic of MG regeneration after LD and NMDA treatment. (B) Inference of activator and repressor function for each individual transcription factor from mulitomic datasets. The y-axis represents the correlation distribution between gene expression and chromVAR score. The top three activator and repressor TFs are shown on the right. (C) Gene regulatory networks of LD and NMDA datasets. (left) Triple regulons model. (right) barplot shows the types of regulons between LD and NMDA datasets. (D) Venn diagram shows the overlap of regulons between LD and NMDA datasets. (E) Enriched gene regulatory networks of LD and NMDA treatment. (left) Enriched Triple regulons model for each condition. (right) barplot shows the number of different types of regulons between LD and NMDA enriched GRNs. (F) An example of stat2 regulons. Color indicates the log2 fold change between the LD and NMDA datasets. (G) Heatmap showing the differentially expressed genes in microglia/macrophages after LD and NMDA treatment. Microglia/macrophage cells are ordered by time points after injury, and an averaged expression level is shown for each timepoint. Color represents mean-centered normalized expression levels. (H) Dotplot showing the key activator TFs for each divergent gene cluster. The size of the dot shows the gene expression ratio, and color indicates the statistical significance of differential expression.

We next identified transcriptional regulatory relationships that were specifically active in either light damaged or NMDA-treated samples. While light damage-specific regulons were predominantly activating – just like injury-induced regulons as a whole – the number of NMDA-specific regulons was much much smaller, and contained roughly equal numbers of activating and repressive regulons (Fig. 4E; Table ST4). Light damage-specific activating regulons were selectively targeted by ascl1a, gli2a, lhx2b, rarab, rarga, rxraa, sox13, and zic2b. In contrast, NMDA-specific activating regulons were heavily targeted by multiple Stat factors, including stat1a, stat1b, and stat2, with these factors also activating one another’s transcription (Fig. 4F; Table ST4). Several factors in this network – including gli3, vax2, and sox11b – were also predicted to directly activate *mmp9* transcription, while atoh7 promotes RGC specification and survival ^36, 37^. Light damage-specific negative regulons were heavily targeted by hmga1a and sox11b, which function primarily as activators in NMDA-treated retina. Conversely, NMDA-specific negative regulons were heavily targeted by rarab and rxraa, in contrast to their function primarily as activators following light damage (Table ST4).

UMAP plots of scRNA-Seq and snRNA-Seq data clearly identified a cell cluster corresponding to microglia and/or macrophages, as indicated selective expression of markers such as *cd45*, *mpeg1.1*, and *csf1ra* (Fig. S3). By analyzing scRNA-Seq data obtained from microglia and/or macrophages in light damaged and NMDA-treated retina, we observed that *il1b*, *stat1a/b*, and *stat2* were all selectively upregulated following NMDA treatment (Fig. 4G), implying that microglial-derived *il1b* might drive the increased Stat signaling seen in NMDA-injured MG. Analysis of activating TFs specific to either injury model showed that while many individual TFs were selectively active in both MG and MGPCs following light damage, the smaller number of activating TFs seen following NMDA treatment was generally specific to individual cell clusters, with Stat factors selectively active in activated MG, and atoh7 active in MGPCs (Fig. 4H).

### Differential organization of gene regulatory networks controlling injury-induced and developmental neurogenesis

To further investigate similarities and differences between gene regulatory networks (GRNs) controlling injury-induced and developmental neurogenesis, we generated multiomic snRNA-Seq/ATAC-Seq from developing zebrafish retina, profiling the full timecourse of developmental neurogenesis. We profiled whole embryo head from three timepoints (24hpf, 30hpf, and 36hpf) and whole retina from six additional timepoints (48hpf, 54hpf, 60hpf, 72hpf, 4dpf, and 5dpf), profiling a total of 52,695 cells (Table ST5). Integrated UMAP analysis of this dataset was able to clearly distinguish separate populations of early- and late-stage RPCs, as well as clusters corresponding to each major retinal cell type (Fig. S6A). We observed that the relative fraction of early-stage RPCs diminished rapidly between 36 and 48hpf, while the fraction of late-stage RPCs peaked at 54hpf, declining rapidly thereafter. As expected, RGCs are the first postmitotic cells to be detected, followed rapidly thereafter by horizontal and amacrine cells and cone photoreceptor precursors. Similar results were observed following reanalysis of previously published scRNA-Seq data obtained from developing zebrafish retina (Fig. S6B) ^38^. The relative fraction of mature cones, rods, bipolars and Muller increases rapidly after 72hpf (Fig. S6C). As is the case in the injured adult retina, each cell cluster in both the multiomic and scRNA-Seq datasets is readily delineated by enriched expression and/or greater chromatin accessibility associated with well-characterized marker genes (Fig. S6D).

We next directly correlated cell-specific patterns of transcription and chromatin accessibility between the developing retina and injured adult retina (Fig. 5A, Fig.S6A). To directly compare GRNs controlling the progression of neurogenesis in injured and developing retina, we integrated the two datasets, projecting them into a common UMAP space (Fig. 5B, Table ST6). We performed a similar analysis for the scRNA-Seq datasets (Figure S6B). Both multiomic and scRNA-Seq data detected distinct clusters corresponding to early and late-stage RPCs, as well as immature precursors corresponding to every major retinal cell type. These cell clusters showed expected changes over developmental time, with early-stage RPCs absent by 60hpf, late-stage RPCs greatly reduced in abundance by 5dpf, and small numbers of RGCs detectable by as early at 24hpf, in line with previous observations ^39^ (Fig. S6C, Table ST5). Small numbers of amacrine and horizontal cells were detected shortly thereafter, with bipolar cell and Muller glia not evident till 60hpf. Although immature photoreceptor precursors were detected as early as 48hpf, large numbers of rods and cones were not observed until 72hfp in the multiome data, and they could not be resolved in the scRNA-Seq data.

**Figure 5:**
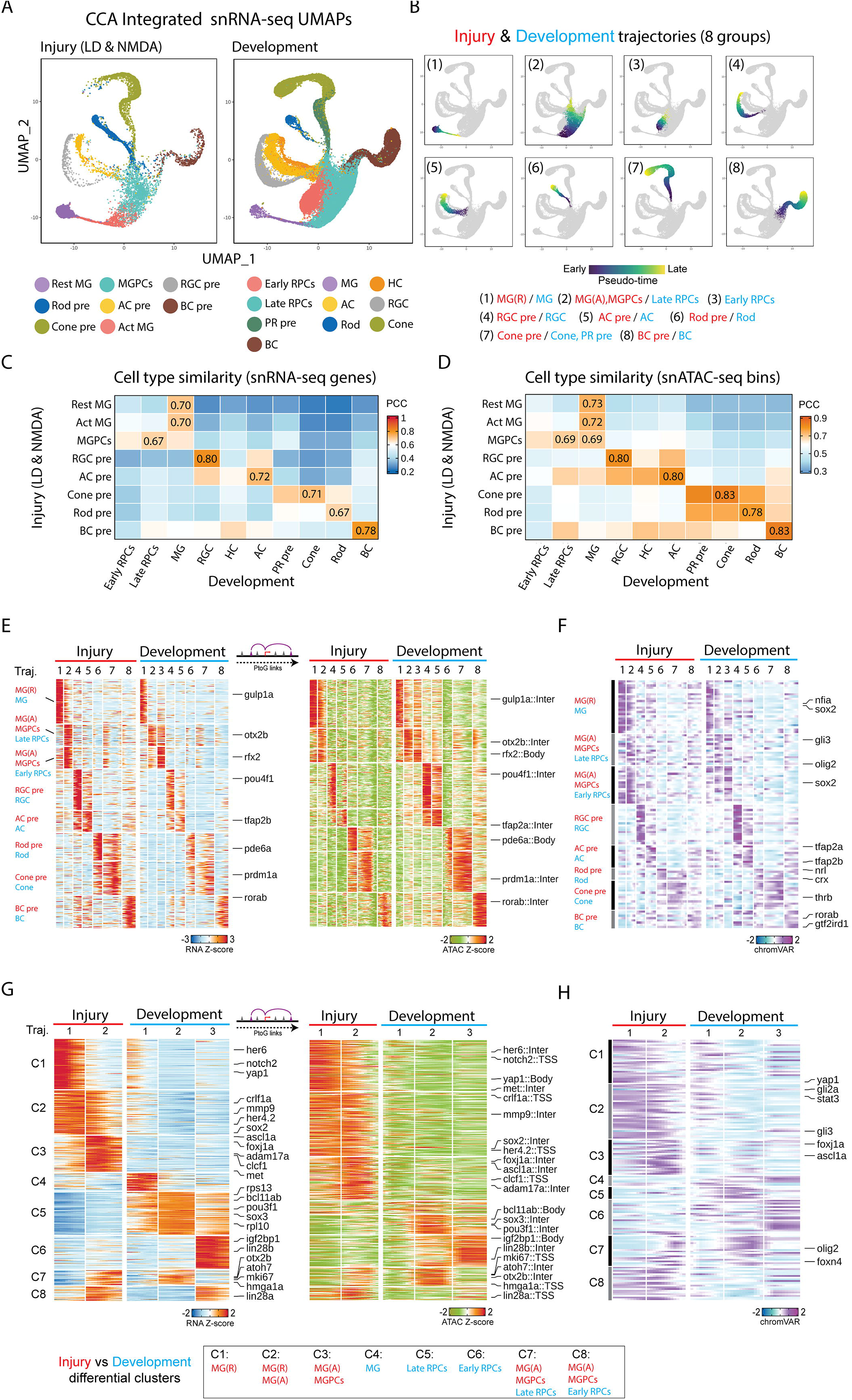
Shared and differential features of MG derived cells between injury and development datasets. (A) Integrated UMAP projection of MG and progenitor neuron cells using injury (left) and development (right) snRNA-seq datasets. Each point (cell) is colored by cell type in each dataset. (B) UMAPs showing the 8 trajectories (groups) constructed from the integrated UMAPs of combined injury and development datasets. Color indicates pseudotime state. The label indicates the cell types included for each trajectory. (C,D) The heatmap displays the Pearson correlations between the cell types from the injury and development datasets using snRNA-seq RNA expression (C) and snATAC-seq bin signals (D). The highest correlation score for each injury cell type is labeled on the heatmap. (E) Heatmap shows the consensus marker genes and their related marker peaks (TSS and enhancer) between injury and development model for each group. (F) Heatmap shows the consensus motifs between injury and development model for each group. (G) Heatmap shows the differential genes and their related differential peaks (TSS and enhancer) between injury and development model for MG groups (group1 and group2). (H) Heatmap shows the differential motifs between injury and development model for MG groups (group1 and group2).

All identified clusters in both datasets expressed well-characterized molecular markers (Fig. S6D). We observed that gene expression patterns in MG from the developmental dataset correlated well between both resting and activated MG from the injury dataset, while MGPCs correlated well with data from late-stage RPCs. We did not, however, observe a distinct early-stage RPC-like cluster in the injury dataset. Immature precursors of each major neuronal cell type likewise correlated most closely with the corresponding cell type present in developing retina, although horizontal cell precursors were absent from the injury dataset (Fig. 5C,D).

To directly compare the process of developmental and injury-induced neurogenesis, we inferred a common trajectory from resting MG to activated MG to MGPCs on the one hand, and from MG to RPCs on the other (Fig. 5E-H). While we observed extensive similarities between gene expression profiles, chromatin accessibility patterns, and TF motif accessibility in both datasets (Fig. 5E,F; Table ST6), we also observed a large number of differentially expressed genes in both trajectories (Fig. 6G,H). Injury-induced trajectory enriched for genes expressed at high levels in mature resting MG (*her6*, *notch2*) and for genes selectively expressed in activated MG (*mmp9*, *clcf1*, *met*, *foxj1a*, *lin28a*) (Fig. 5E, Table ST6). Gene Ontology and KEGG analysis shows that genes associated with cell adhesion, neurogenesis, and FoxO and VEGF signaling are enriched in the injury dataset, while genes controlling cell cycle, protein synthesis and splicing were enriched in the developmental dataset (Fig. S4C,D). We likewise observed major differences in both differential chromatin accessibility (Fig. 5G) and TF motifs (Fig. 5H) between the injury and developmental datasets. Notably, we observe that the promoters of many genes selectively expressed in resting and/or activated MG – including *her6*, *met*, *mmp9*, *lin28a*, and *foxj1a* – are not accessible in RPCs in the developing retina (Fig. 5G). We likewise observe that target motifs for some of these same factors, including stat3 and foxj1a, are likewise accessible only in the combined injury dataset (Fig. 5H).

**Figure 6:**
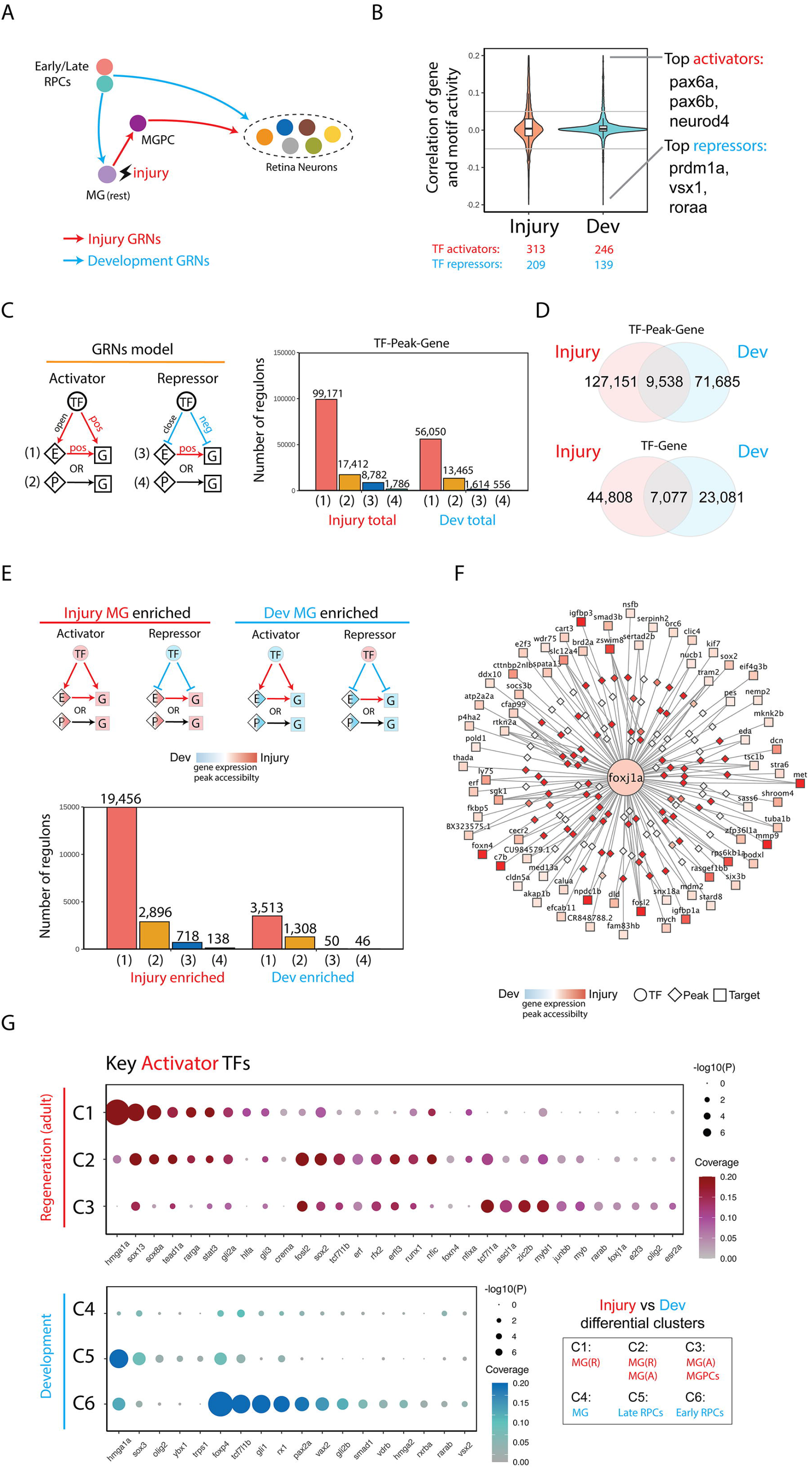
Transcription factors controlling differential expression genes in MGPC in injured retina and progenitor cells in developing retina. (A) Schematic of MG regeneration between injury and development. (B) Inference of activator and repressor function for each individual transcription factor from mulitomic datasets. The y-axis represents the correlation distribution between gene expression and chromVAR score. The top three activator and repressor TFs are shown on the right. (C) Gene regulatory networks of injury and development datasets. (left) Triple regulons model. (right) barplot shows the number of types of regulons. (D) Venn diagram shows the overlap of regulons between injury and development GRNs. (E) Enriched gene regulatory networks of injury and development. (Left) Enriched Triple regulons model for each condition. (Right) barplot shows the number of different types of regulons between injury and development enriched GRNs. (F) An example of foxj1a regulons. Color indicates the log2 fold change between injury and developmental datasets. (G) Dotplot showing key activator TFs for each divergent gene cluster. The size of the dot showing the gene ratio and color indicates the significance of regulation.

### Identification of gene regulatory networks differentially regulating neurogenesis in MGPCs and RPCs

Having observed extensive similarities and differences in both gene expression and chromatin accessibility over the course of neurogenesis in MGPCs and RPCs, we next reconstructed and directly compared dynamic GRNs in both cell types using the same methods we previously used to compare GRN in NMDA and light-damaged MG and MGPCs (Fig. 6A). The aggregated light damage and NMDA Injury-regulated GRNs consisted of 522 unique TF-Motif pairs, with developmental TFs comprising 385 of these, with a roughly 60:40 ratio of activators to repressors (Fig. 6B). Analysis of injury and development-associated regulons showed that 91.6% of injury-associated and 97.0% of development-associated regulons were activating (Fig. 6C). 11.7% of development-specific TF-peak-gene interactions and 23.5% of TF-gene interactions overlap with the injury-specific dataset (Fig. 6D). Analysis of injury and development-specific regulons reveals that these are likewise dominated by activating relationships, with more than five times as many total injury-specific regulatory relationships than developmental-specific regulatory relationships (Fig. 6E).

Development-enriched activating TFs include hmga1a, foxp4, pax2a, rx1, and sox3. Injury-enriched activating TFs include ascl1a, rfx2, sox2, sox13, stat3, tead1a, and zic2b (Fig. 5G,H; Table SX). An example of this category of key injury-induced regulatory TF is shown for *foxj1a*, which is both selectively expressed in the regenerating retina (Fig. 5G,H; Fig. 6F). We observe that *foxj1a* directly regulates several genes that regulate MGPC formation, such as *mmp9, foxn4,* and *sox2* ^18, 33, 40^. Notably, *foxj1a* is also more strongly induced following light damage than NMDA treatment (Fig. 2G), indicating that it may also differentially regulate injury-induced neurogenesis in these two models. We observe that many of the injury-enriched activating TFs are strongly and selectively expressed in activated MG, which are absent from the developmental dataset, while development-enriched TFs are predominantly active in early-stage RPCs, which are likewise lacking from the injury-induced dataset (Fig. 6G).

### *Foxj1a* is necessary for neuronal regeneration of the damaged adult retina, but it is not required for retinal neurogenesis

To determine if any of these candidate injury-induced TFs are required for neuronal regeneration, but not in retinal development, we tested *foxj1a* (Fig. 5G,H; 6F). *foxj1a* is expressed much higher during retinal regeneration than retinal development (Figure 7A,B). To test the role of *foxj1a* in regeneration, morpholinos were intravitreally injected and electroporated into either NMDA-damaged or light-damaged retinas at the start of damage ^41^. As controls, we used the Standard Control morpholino, which is not complementary to any known sequence in the zebrafish genome (GeneTools, LLC), and the *pcna* (proliferating cell nuclear antigen) morpholino, which is necessary for MG and MGPC proliferation during retinal regeneration ^42^. As expected, the *pcna* morphant retina possessed significantly fewer PCNA-positive proliferating MGPCs at 72 hours following either NMDA damage (Fig. 7C-E) or light damage (Fig. 7F-H). We likewise observed significantly reduced levels of PCNA+ cells in the ONL and INL of both NMDA-treated and light damaged retinas following treatment with the *foxj1a* morphant. However, while the *pcna* and *foxj1a* morphants were equally effective at inhibiting proliferation in the NMDA-treated retina (Fig. 7D,E), the *foxj1a* morphant was less effective than the *pcna* morphant at doing so in light-damaged retinas (Fig. 7F,G).

**Figure 7:**
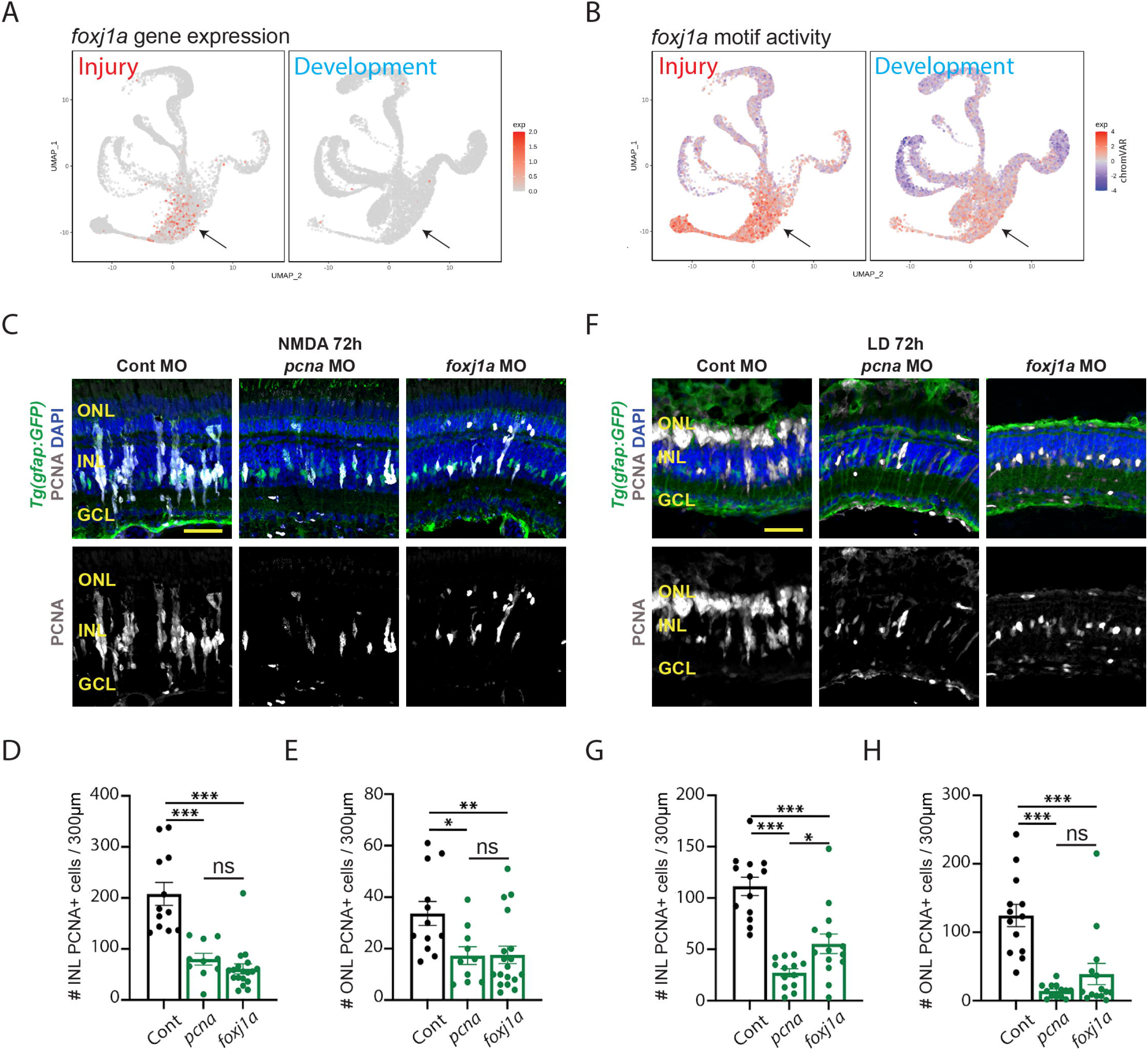
Foxj1a is required for MGPC proliferation. (A). UMAPs showing the gene expression pattern of foxj1a between injury (left) and development model (right). (B). UMAPs showing the chromVAR motif activity of foxj1a between injury (left) and development model (right). (C) Tg(*gfap:GFP*) retinas electroporated with either Standard Control morpholino (Cont MO), *pcna* MO, or *foxj1a* MO were isolated 72 hours after NMDA injection and immunostained for PCNA, GFP, and counterstained with DAPI. (D) Quantification of the number of PCNA-labeled cells in the INL. (E) Quantification of the number of PCNA-labeled cells in the ONL. (F) Tg(*gfap:GFP*) retinas electroporated with either Cont MO, *pcna* MO, or *foxj1a* MO were isolated 72 hours after starting constant light and immunostained for PCNA, GFP, and DAPI. (G) Quantification of the number of PCNA-labeled cells in the INL. (H) Quantification of the number of PCNA-labeled cells in the ONL. Scale bars in A and D are 20μm.

Furthermore, at 36 hours following injury, while the *pcna* morphant was effective at inhibiting proliferation of INL cells (which correspond to proliferating MG at this stage) in both NMDA-treated and light-damaged retina, the *foxj1a* morphant only reduced proliferation in NMDA-treated retinas at this stage, and had no significant effect in light-damaged retina (Fig. S7A-F). At this time, proliferating ONL cells represent rod precursors ^43^, which are not derived from injury-induced MGPCs and do not take up either morpholino, and as expected showed no change in either morphant (Fig. S7A-F). Loss of function of foxj1a therefore preferentially affected MG proliferation in NMDA-treated retina relative to light-damaged retina. All cell counts are listed in Table ST1.

To determine if *foxj1a* is required for developmental retinal neurogenesis, wild-type embryos were injected at the 1-4 cell stage with either uninjected or injected with the Standard Control morpholino, *mmp9* morpholino, or the *foxj1a* morpholino. At 96 hpf, all the embryos appeared normal except for the *foxj1a* morphants, which possessed significantly curled body axes (Fig. S7G), which is consistent with the previously published *foxj1a* mutant phenotype ^44^. At 96 hpf, we examined retinal organization by immunostaining (Fig. S7G). Both the uninjected and the Standard Control morphant retinas possessed normal lamination and wild-type staining for rod photoreceptors, HuC/D (amacrine and ganglion cells), and GFAP (MG; Fig. S7G). While the *foxj1a* morphant retinas possessed normal lamination and all the antibodies stained the expected retinal layers, the overall maturation of the retina appeared somewhat delayed relative to the control retinas (Fig. S7G). It is unclear if this delayed retinal development seen in *foxj1a* morphants is due to its previously documented widespread early developmental function ^44^, or is mediated by disrupted function of the very small number of *foxj1a*-expressing cells in late retinal progenitor cells. These findings demonstrate a selective role for foxj1a in regulating injury-induced neurogenesis in zebrafish retina.

## DISCUSSION

In this study, we demonstrate that different retinal damage models induce a similar cellular response by which the MG reprogram and divide asymmetrically to produce MG-derived progenitor cells that continues to proliferate and differentiate into retinal neurons. However, there are distinct differences in the gene expression profiles and TF networks between these two retinal damage models. For example, *mmp9*, which is expressed at a higher level in NMDA-damaged retinas relative to light-damaged retinas, represents the first identified gene that selectively regulates generation of set of MGPC-derived retinal cell types (amacrine and ganglion cell cells) relative to another (photoreceptors). Further, while these two damage models produce multipotent MGPCs that are transcriptionally similar to late retinal progenitors, there are several differences in both gene expression and transcriptional regulatory networks between MGPCs and bona fide late-stage progenitors from developing retina. This is exemplified by the identification of specific genes, such as *foxj1a*, that significantly repress retinal regeneration while only modestly delaying retinal development. Taken together, these results reveal that while there are similarities in the gene expression and TF networks between different damage models and between regenerating and developing retinas, key differences exist that highlight the finding that regeneration is not a direct recapitulation of development at the level of gene regulation, and that different gene and TF networks are required for the proper regeneration of retinal neurons in different damage environments.

Our comparative analysis of MGPCs following light damage and NMDA injury reveals several common and often unexpected features. We observe that MGPCs mostly produce rod and cone photoreceptors, amacrines, and RGCs regardless of the injury model, while generating few bipolar and no horizontal cells. MGPCs also show inherent biases in neurogenic competence that are determined by the specific neuronal cell types lost to injury. Light damage preferentially induces generation of rods and cones, while NMDA preferentially generates RGCs. However, there is extensive generation of most retinal neuronal cell types from MGPCs in both injury models.

### Postmitotic elimination of MGPC-derived neurons appears to be minimal

These biases in neurogenic competence are likely the result of differential transcriptional response to extrinsic signals. Light damage induces substantially higher activity of neurogenic bHLH factor *ascl1a*, while NMDA injury triggers a more typical inflammatory response, marked by particularly high expression and activation of *stat1/2* genes. MGPCs show substantially higher expression of several TFs that promote formation and/or survival of RGCs, such as *atoh7*, and these likely play an important role in restricting developmental competence. While light damage induces higher levels of expression of *foxj1a* relative to NMDA-treated retinas (Fig. 2G), *foxj1a* morphants show a less dramatic reduction in MGPC proliferation in light-damaged retinas (Fig. 7). This may result from there simply being more functional foxj1a protein present in light-damaged retinas following morpholino treatment, or else may point to unexpected regulatory functions of this transcription factor, which await further functional investigation. Although foxj1a is well-characterized as a master regulator of the development of multiciliated ependymal cells ^45^, these are not present in either the developing or regenerating retina, and no evidence for selective expression of ciliary genes is seen in the regenerating retina (Table ST6). The recent finding that Foxj1 regulates mammalian cortical cell fate specification independent of any role in ciliogenesis ^46^ identifies new potential mechanisms of action in the context of MGPC proliferation.

Secreted factors such as Mmp9 likewise play an important role in selectively controlling generation of specific cell types. Mmp9 has been previously shown to inhibit generation of MGPC-derived photoreceptor precursors following light damage ^33^. While we replicate this observation here, we also observe a stronger and more selective effect on inhibiting generation of amacrine and ganglion cells following both light damage and NMDA injury. The fact that *mmp9* expression is enriched in MGPCs in the NMDA-treated retina, which are biased towards generating ganglion cells, underscores the importance of precise control of injury-induced neurogenesis through active negative regulation. The precise mechanism of action of mmp9 in this context remains unclear. Mmp9 is a secreted protease that cleaves components of the extracellular matrix such as collagen and laminin ^47^, as well as multiple chemokines ^48^ and lipid binding proteins ^49^, and both MG and microglia differentially express multiple genes in all these functional categories following NMDA treatment and light damage (Table ST3). Most notably, mmp9 also controls maturation of il1b ^50^, which is selectively upregulated in microglia following NMDA injury (Fig. 4G). Further experiments are required to determine the mechanism by which mmp9 selectively inhibits generation of inner retinal neurons.

It was previously described that zebrafish retinal development begins with the commitment of retinal ganglion cells, followed by cells in the INL, then the cone photoreceptors and ultimately the rod photoreceptors and Muller glia ^39^. Our snRNA-Seq data is consistent with this temporal pattern (Table ST5). However, our data also suggests some additional refinement of this pattern. For the INL cells, amacrine cells appear to be generated immediately following the ganglion cells. The bipolar cells and horizontal cells are next and, as described previously ^39^, we then observe generation of cone photoreceptors, followed by the rod photoreceptors. Transcriptionally distinct MG precursors are generated between the cone and rod photoreceptors. This suggests a model of regeneration where the MG reprogram and produce MGPCs that are very similar to late-stage RPCs, which then first generate ganglion cells, followed by INL neurons, and then cone and rod photoreceptors. We propose that this results in the differentiation of all retinal neuronal classes, regardless of the neuronal classes lost due to damage. However, if the early differentiating neuronal classes experienced little, or no, cell loss, then the majority of the MGPCs are retained and differentiate into photoreceptors. In contrast, if the damage results in primarily inner retinal neuronal classes being lost (i.e. NMDA damage), then the majority of the MGPCs must differentiate into these lost neuronal cell types and only a subset of the MGPCs are retained to differentiate into cone and rod photoreceptors. This model is in addition to the damage-specific transcriptional changes that we observed, which can further refine the commitment of the MGPCs into the differentiated neuronal ratios that were observed.

While MGPCs closely resemble RPCs at the molecular level, a number of major differences are evident that likely correspond to known differences in proliferative and/or developmental competence. RPCs show a distinct early-stage state that is not observed in MGPCs. This does not appear to strictly correspond to an early state of developmental competence, as has been reported in mammals ^51–53^, since early-born cell types such as RGCs, horizontal cells and cone photoreceptors are still being generated at 48 hpf, when RPCs have adopted a late-stage profile. These differences between RPCs and MGPCs may instead correspond to differences in proliferative competence, as symmetric non-neurogenic cell divisions predominate in early-stage zebrafish but are not detected in MGPCs ^13, 54^. Furthermore, while RGCs are efficiently generated by MGPCs, they are generated at much higher relative levels during early stages of developmental neurogenesis, which likely reflects that fact that a large majority will be eventually eliminated by apoptotic cell death ^55^. The relatively weaker expression of RGC-promoting factors such as *sox11a* in MGPCs may mediate these differences in RGC generation, while the absence of horizontal cell-promoting TFs such as *onecut1* and *pou2f2a* likely underpins the lack of competence to generate horizontal cells seen in MGPCs in these injury models. It remains to be determined whether selective conditional ablation of either cell type ^56^ might activate expression of these genes in MGPCs. Furthermore, since robust methods of inducing MG-derived neurogenesis have now been developed in mammals ^38, 57–59^, these findings raise the possibility that different modes of injury might also differentially regulate generation of specific MG-derived neuronal subtypes in this context.

## AUTHOR CONTRIBUTIONS

SB, JQ, and DRH conceived and supervised the study. PL analyzed all the mutiomic data generated, while YZ analyzed all the scRNA-Seq data. MI and DS generated and analyzed all of the NMDA-treated and light-damaged data, and conducted all functional studies of *mmp9* and *foxj1a* in adult retina, with assistance from PB. TH generated all the scRNA-Seq, snRNA-Seq, and scATAC-Seq data, with assistance from IP. LJC conducted all analysis of *foxj1a* morphants in the early embryo. MN, NJS, and PFH generated and provided the *mmp9* mutants. PL, MI, DS, YZ, JQ, DRH, and SB drafted the manuscript. All authors edited the manuscript.

## DATA AVAILABILITY

All scRNA-Seq, snRNA-Seq, and scATAC-Seq data have been deposited in GEO (GSE239410).

## ACKNOWLEDGEMENTS

This work was supported by the Milky Way Research Foundation (to SB, DH, and JQ), and by NIH grants, R01EY007060, R21EY034182, P30EY007003, R01EY034493 and an unrestricted grant from the Research to Prevent Blindness (to PFH). We thank Jeremy Nathans, Alex Kolodkin, Jeff Mumm, and Andrew Fischer for comments on the manuscript.

## METHODS

### Zebrafish husbandry

Adult *albino^b^*^4^*^/b^*^4, 60, 61^, *mmp9^mi^*^5003^ mutant fish ^33^, and transgenic *albino Tg(gfap:EGFP)^nt^*^11, 19^ zebrafish used in this study were bred and maintained in the Freimann Life Science Center Zebrafish Facility at the University of Notre Dame in accordance with standard operating policies and procedures ^2, 33^. The fish were kept at a temperature of approximately 26.5°C and 30% humidity, with a 14-hour light, 10-hour dark cycle ^62^. *Tg(gfap:EGFP)^nt^*^11^ was used to label the MG by expressing GFP ^19^. All the zebrafish used in this study were adults between 6 and 12 months old or embryos. Prior to any experiments, the fish were anesthetized or euthanized using 1:1000 or 1:500 2-phenoxyethanol (Sigma), respectively.

### Retinal damage and EdU labeling

Phototoxic treatment that causes photoreceptor ablation was used to initiate a regenerative response in the zebrafish retina ^2, 63^. All fish subjected to light damage were dark-adapted for two weeks and then placed adjacent to intense fluorescent lights for 96 hours. The temperature of the fish tanks was regularly monitored and maintained at 34°C. After 96 hours, the fish were returned to the animal facility with a normal light/dark cycle and maintained for 7, 14, or 21 days recovery (DR).

N-methyl-D-aspartate (NMDA, Sigma M3262) was administered to normal light/dark cycled fish via an intravitreal injection of 0.3µL 100mM NMDA solution in sterile PBS with a Hamilton syringe (Hamilton, 2.5µL Model 62 RN SYR, 87942), followed by placing the tanks in a dark incubator at 33°C for 96 hours, after which the tanks were moved back into a normal light/dark cycle for 7 or 14 days recovery.

EdU (5-Ethynyl-2′-deoxyuridine, Life technologies A10044) was administered as previously described ^63^. Fish were anesthetized using 1:1000 2-phenoxyethanol and immobilized under a paper towel in a petri dish. Then, using an insulin syringe equipped with a 30-gauge needle, approximately 20 µL 1mg/mL EdU in water was injected intraperitoneally into each zebrafish at 60, 72, 84, 96 and 108 hours after damage. The retinal tissues were harvested at either 3, 7, 14, or 21 days recovery.

### *In vivo* morpholino-mediated gene knockdown

Conditional adult retinal morphants were generated essentially as described previously ^41, 42^. Prior to light treatment, fish were anesthetized in 1:1000 2-phenoxyethanol, the outer cornea membrane of the left-eye was removed with Dumont #5 forceps, an incision was then made though the inner cornea in the ventral-caudal aspect of the pupillary margin using a #11 scalpel, and 0.2-0.3µL of morpholinos (Gene Tools, LLC) at a concentration of 1mM were injected into the posterior segment using a 2.5µL Hamilton syringe. Morpholinos that do not target any known zebrafish genes (Standard Control), as well as one targeting *pcna*, which blocks MG proliferation, were used as negative and positive controls, respectively. Morpholinos were electroporated into the retina using platinum electrode tweezers with a CUY21EDIT Square Wave Electroporator (Nepa Gene Company Ltd.), delivering two 50 msec pulses, separated by a 1 second pause, at 0.75 Volts, resulting in the current of 0.05 Amperes. The electrodes were used to prolapse the eye out of the socket, with the anode in contact with the cornea, and the cathode positioned 2mm behind the dorsal part of the eye. Successful electroporation was confirmed by the presence of lissamine fluorescence within the retinal cells using microscopy. For the embryos conditional morpholinos mediated knock-down experiments, the morpholinos were diluted to 1 mg/mL in 1x Danieau buffer. Wild-type or *Tg(gfap:EGFP)^nt^*^11^ embryos were injected with 5 ng of morpholinos at the one-cell stage ^64^.

Morpholino sequences used were: Standard Control: 5’ CCTCTTACCTCAGTTACAATTTATA 3’ (Gene Tools, LLC), anti-*pcna:* 5’ TGAACCAGACGTGCCTCAAACATTG 3*’* ^42^, anti-*foxj1a:* 5’ CATGGAACTCATGGAGAGCATGGTC 3’ ^65^, and anti*-mmp9:* 5’ ACTCCAAGTCTCATTTTGAGTCGCA 3’ (Gene Tools, LLC).

### Tissue fixation and cryosectioning

Tissues for immunocytochemical analyses were collected at specified time points by euthanizing zebrafish in 1:500 solution of 2-phenoxyethanol, followed by ocular enucleation using #5/45 Dumont forceps (Supplier). The eyes were floated in 1x Phosphate-Buffered Saline (PBS) solution, and a circular opening, approximately the size of the pupil, was made in the cornea using micro-scissors, creating an eye cup. The eye cups were fixed at room temperature (RT) in 4% paraformaldehyde, using a pH shift technique to preserve the cytoskeletal structure and avoid the cellular swelling that is typically associated with standard paraformaldehyde fixation at 4°C ^66^. Briefly, the tissues were fixed at RT for 5 minutes in 80mM HEPES, 2mM EGTA, 2mM MgCl_2_· 6H_2_O, 4% PFA, pH 6.8-7.2, followed by 30 minutes in 100mM Na_2_B_4_O_7_· 10H_2_O, 1mM MgCl_2_· 6H_2_O, 4% PFA, pH 11. The eyes were washed in 1x PBS at RT and dehydrated in a 15% sucrose solution at 4°C overnight. The eyes were then placed in a 2:1 solution of PolyFreeze Tissue Freezing Medium (TFM) (Sigma-Aldrich) and 30% sucrose for 24 hours at 4°C. Finally, the eyes were arranged in the dorsal-ventral plane in 100% TFM, frozen, and stored at −80°C. Frozen tissue blocks were sectioned using a Thermo Scientific Microm HM550 Cryostat at −23°C, producing 14µM serial sections that included the optic nerve and both the dorsal and ventral crescent. Retinal sections were collected on SuperFrost glass slides (VWR 48311-703), dried for 30 minutes on a slide warmer at 50°C, and stored at −80°C for future immunohistochemical staining.

### Immunohistochemistry

Retinal sections were processed for immunohistochemistry as previously described ^2, 63^. When anti-PCNA primary antibodies were used, an extra heat-induced antigen retrieval step was performed. Prior to blocking, slides were immersed in a 10mM Sodium Citrate, 0.05% Tween 20, pH 6 solution, and heated for 25 seconds in a microwave at maximum power to nearly reach the boiling point. The slides were heated for another 7 minutes using the microwave at 10% power in periodic 20-25 second bursts to maintain near-boiling temperature. The slides were cooled for 30 minutes at RT and washed 2×5 minutes in 1xPBS before proceeding to the blocking step.

The following primary antibodies were used: mouse anti-PCNA monoclonal antibody (Sigma P8825, 1:500 dilution), rabbit anti-GFAP (Dako Z0334, 1:300 dilution), mouse anti-HuC/D monoclonal antibody (Invitrogen A21271, 1:300 dilution), mouse anti-4c12 monoclonal antibody (gift from Dr. Fadool, Florida State University, 1:200 dilution), rabbit anti-PKCa (Sigma Life Science P4334, 1:300 dilution), rabbit anti-UV cone opsin ^67^, 1:1000 dilution), rabbit anti-green cone opsin ^67^, 1:500 dilution), mouse anti-Zpr1 monoclonal antibody (ZIRC, 1:200) rabbit anti-Lcp1 (GeneTex GTX134697, 1:500). Secondary antibodies (diluted 1:500) included: goat anti-mouse 488 (Life Technologies A11029), goat anti-mouse 594 (Life Technologies A11032), goat anti-mouse 647 (Life Technologies A21236), goat anti-rabbit 488 (Life Technologies A11034), goat anti-rabbit 594 (Life Technologies A11037), goat anti-chicken 488 (Life Technologies A11039).

### Confocal microscopy and quantification

A Nikon A1R HD inverted confocal microscope at the University of Notre Dame Integrated Imaging Facility was used to capture serial optical sections of retinal slices using 40x and 60x oil-immersion objectives. Z-stacks of approximately 10µM from the central dorsal retinal tissue were captured using NIS-Elements (RRID:SCR_014329). The resulting images were analyzed as either single optical sections or maximum z-stack projections (where appropriate) using ImageJ/FIJI ^68^. For the colocalization experiments, single optical sections were used. We increased the saturation of the Gfap and 4c12 channels to increase the cell body staining to determine colocalization with EdU.

### Statistical analysis

The resulting data were analyzed using GraphPad Prism version 9. Student’s t-test for single pairwise comparisons with control or one-way or two-way analysis of variance followed by Dunnett’s Bonferroni’s post hoc test for multiple comparisons. The statistical significance is indicated in each graph as follows: *p ≤ 0.05, ** for p ≤ 0.01, *** for p ≤ 0.001, or ns for not significant difference.

### ScRNA-Seq of methanol-fixed retinal cells

Zebrafish retinas (4-6 retinas from 4-6 fish) were collected at different timepoints after NMDA and LD treatments. Retinal cells were dissociated and fixed in −20^0^ C methanol and stored in −80^0^ C freezer until used. Fixed cells were recovered and subjected to 10x Genomics Single Cell system as described previously ^38^. Libraries were prepared and sequenced on Illumina NovaSeq at ∼500 million reads per library

### Multiome analysis of frozen retinal tissues

Zebrafish retinas (4-6 retinas from 4-6 fish) were collected at different timepoints after NMDA and LD treatments, and flash-frozen in dry ice for ∼15min before being transferred to a −80^0^ C freezer for storage. Nuclei were extracted from frozen retinal tissues according to 10xMultiome ATAC + Gene Expression (GEX) protocol (CGOOO338). Briefly, frozen retinal tissues were lysed in ice-cold 500ml of 0.1X Lysis buffer using a pestle and incubated on ice for 6 min totally. Nuclei were centrifuged, washed 3 times and resuspended in 10xMultiome nuclei buffer at a concentration of ∼3000-5000 nuclei/ml. Nuclei (∼15k) then were loaded onto 10x Genomic Chromium Controller, with a target number of ∼10K nuclei per sample. RNA and ATAC libraries were prepared according to the 10xMultiome ATAC + Gene Expression protocol and subjected for Illumina NovaSeq sequencing at ∼500 million reads per library

### Single-cell muti-omics analysis Preprocessing

The multi-omic sequencing files were processed for demultiplexing and analyzed using Cell Ranger ARC v2.0. The genes were mapped and referenced using the zebrafish reference genome DanRer11. To address potential ambient RNA contamination in the cell-by-gene expression matrix, we employed scAR ^69^ module from scvi-tools ^70^ with default parameters for further ambient RNA cleaning for each sample.

### Filtering barcode doublets and low-quality cells for each sample

The fragments files were first converted to ArchR ^71^ objects and the cell-by-genes matrices were converted to Seurat v3 objects ^72^. The doublet cells were identified using the solo module^73^ of scvi-tools with the cleaned cell-by-gene RNA matrix and the ArchR package with the cell-by-bin ATAC matrix. If the Solo doublet score of a cell was greater than zero or the ArchR doublet enrichment score was greater than 2, the doublet cells were filtered out. Furthermore, cells with low RNA counts (nCount_RNA < 500) and low ATAC counts (nCount_ATAC < 1000), as well as cells with high RNA counts (> 50,000) and high ATAC counts (> 100,000), were further filtered out as low-quality cells. Additionally, cells were also filtered based on their transcription start site (TSS) enrichment (< 4).

### Clustering, visualization, and identification of cell types

The clustering analysis was conducted on the combined dimensional space obtained from both the RNA and ATAC single-cell matrices. Initially, the RNA expression matrix was incorporated into the ArchR project using the “addGeneExpressionMatrix” function. Subsequently, dimension reduction was separately applied to the cell-by-bin matrix and the cell-by-gene matrix within the ArchR project using the “addIterativeLSI” function. The resulting reduced dimensions were then merged into a new dimension called “LSI_Combined” using the “addCombinedDims” function. Finally, clustering was performed using the “addClusters” function with a resolution of 1.5, taking into account the “LSI_Combined” dimensions. To visualize single cells, the UMAP embeddings were calculated using the “addUMAP” function in ArchR with the first 10 “LSI_Combined” dimensions.

To infer cell types, the existing zebrafish scRNA-seq data ^38^) were utilized for interpreting our snRNA-seq cell types using the CCA (canonical correlation analysis) integration method in the Seurat package ^74^. Firstly, the zebrafish scRNA-seq data were downloaded and converted into Seurat objects. Secondly, for each snRNA-seq sample, the anchors between them were identified using the ‘transfer.anchors’ function, and the ‘TransferData’ function was used to obtain the cell type prediction results for each cell. Cells with a prediction score < 0.5 were further filtered out, and each cluster was annotated based on their predicted cell types. To further identify precursor cell clusters, the clusters in each major cell type between the control and injury datasets were compared. Clusters that existed in the injury dataset but not in the control dataset were selected as precursor cell clusters and were further confirmed using specific precursor cell markers.

### Generating fixed-width and non-overlapping peaks

Fixed-width 501 bp peaks were called for each cell type in both the injury and development datasets by MACS2 ^75^ in ArchR package. Subsequently, these peaks were merged using an iterative method to retain the most significant peak among overlapping peak sets. The function “addReproduciblePeakSet” was utilized for this calling and merging process. Following that, the peak matrix was generated for each dataset using the function “addPeakMatrix” with the following parameters: ceiling = 4 and binarize = FALSE.

### Single-cell RNA-seq data analysis Preprocessing

The scRNA sequencing files were processed for demultiplexing and analyzed using Cell Ranger. The genes were mapped and referenced using the zebrafish reference genome DanRer11. To address potential ambient RNA contamination in the scRNA-seq expression matrix, we employed SoupX ^76^ with default parameters for further ambient RNA cleaning for each sample.

### Quality control of scRNA-Seq data for each sample

The cell-by-genes count matrices were converted to Seurat v3 objects ^72^. Cells with RNA counts less than 500 or greater than 50,000 were filtered out as low-quality cells. Additionally, cells with a mitochondrial fraction of greater than 15% were also removed. Doublet cells were identified using the Scrublet Python package ^77^.

### Clustering, visualization, and cell type annotation of scRNA-Seq data

The control, LD and NMDA samples were first integrated using the Seurat integration functions (SelectIntegrationFeatures, FindIntegrationAnchors and IntegrateData). Data scaling, dimensional reduction and clustering were then performed on the integrated injury dataset using the standard Seurat pipeline. For visualization, combined UMAP were generated using the first 20 dimensions. We used the same methods to infer cell types as we did for single-cell multiomics data. Briefly, the existing zebrafish scRNA-seq data ^38^) were utilized for interpreting our scRNA-seq cell types using the CCA (canonical correlation analysis) integration method in the Seurat package.

### Integration of injury and development single-cell RNA-seq data

The similarity between the injury and development datasets was assessed by integrating them based on their single-cell expression profiles using the reciprocal PCA workflow from Seurat. The integration process was performed with a strength parameter of k.anchor = 5. Subsequently, integrated UMAPs were generated using the first 1 to 15 dimensions derived from the integrated dimensions. The cell type annotations on the new integrated dataset remained the same as those in the separate injury and development datasets.

### Trajectory inference and pseudotime analysis

Slingshot software ^78^ was utilized to infer pseudotime based on the UMAP coordinates, which were filtered by selecting cells involved in the specific biological process under investigation.

For the injury datasets, the LD and NMDA combined UMAP coordinates were first separated based on the cells from the 6 groups: MG (Rest MG, Activated MG and MGPCs); RGC pre; AC pre; BC pre; Cone pre; and Rod pre. Then, Slingshot was applied to infer trajectories using the ‘getLineages’ and ‘getCurves’ functions. In the Rest MG to MGPCs group, the cell cluster overlapping with “Rest MG” was treated as the “root cluster”. For the other groups, the closest cell cluster to “MGPCs” was treated as the “root cluster”.

For the combined injury and development datasets, Slingshot was run to construct MG-related trajectories using the merged scRNA-seq UMAP coordinates. The cell cluster overlapping with “Rest MG” in the injury dataset and “MG” in the development dataset was treated as the “root cluster” for Slingshot.

Subsequently, the ‘slingPseudotime’ function was utilized to calculate the pseudotime state for each cell. Finally, the pseudotime was merged into 20-50 bins for each trajectory, and the average gene expression, average accessibility level, and average motif activity score were calculated for each bin. The scores for each bin were further smoothed and scaled for visualization.

### Identification of Differential and Consensus Genes, Peaks, and Motif Activities

The consensus (CEG) and differential genes (DEG) were identified using the ambient-RNA-cleaned cell-by-gene count matrix for both scRNA-seq and multi-omic datasets. The peaks were annotated with their predicted target genes, as described in the GRNs construction methods section. To further remove potential ambient RNA contamination from DEGs, the marker genes of mature retina neurons (Rod, Cone, RGC, AC, HC, BC, Microglia) were identified using control samples with the FindMarkers function: log2FC > 2 and adj-pvalue < 0.05. The DEG list of MG cell groups (MG, Act MG, MGPCs) in the downstream analysis excluded these mature marker genes. In the motif analysis, motifs from both cis-BP and Transfac database (2018) were collected. For each TF, the corresponding motifs were filtered based on the correlation between TF expression and motif activity (chromVAR) score, which described in the GRNs construction methods section.

To identify the DEGs of MG cell groups between the LD and NMDA treatment (or between injury and development) datasets, the “findMarkers” function in the Seurat package was employed. For the LD vs NMDA comparison, the DEGs were defined as abs(log2FC) > 0.25 and adj-pvalue < 0.05. For the injury vs development comparison, the DEGs were defined as abs(log2FC) > 0.5 and adj-pvalue < 0.05. Finally, all the DEGs are merged and clustered using k-means method by their expression profile along the trajectories of MG groups. (Figure 2G and Figure 5G)

To identify the CEGs between different conditions, the “findAllMarkers” function was used to determine enriched genes for each trajectory group compared to other groups. CEGs were identified based on the following criteria: a log2FC greater than 0.5 and an adjusted p-value less than 0.05 in both the LD and NMDA datasets (or in both injury and development datasets).

The differential accessible regions (DARs) between LD and NMDA datasets (or between injury and development) were identified using the “getMarkerFeatures” function in ArchR to measure the peak differences between LD and NMDA. DARs were identified for each cell group in the injury datasets based on the criteria of having an absolute log2 fold change greater than 0.25 and a p-value less than 0.05. Additionally, DARs were further filtered if their predicted target genes are not DEGs.

To identify the consensus lineage-specific peaks (CARs) between LD and NMDA datasets (or between injury and development), peaks were separately identified for each group compared to other groups in each condition. CARs were considered significant if they exhibited a log2 fold change greater than 0.25 and an pvalue less than 0.05. Furthermore, CARs were excluded if they were not related with the CEGs.

The differential motifs (DMs) between the LD and NMDA treatment datasets (or between injury and development) were identified using the “FindMarkers” function from the “Signac2” package. The cell-by-motif z-score matrix from chromVAR was converted into a Seurat object. The “LR” test method was utilized to determine the differential motif accessibility. Motifs with an absolute log2 fold change (log2FC) greater than 0.25 and an adjusted p-value less than 0.05 were classified as DMs. Furthermore, DMs were excluded if they were found to be inconsistent with the list of differentially expressed genes (DEGs) for each cell type. In cases where multiple DMs corresponded to a DEG in the list, only the DM with the highest correlation with its corresponding DEG was retained.

The consensus motifs (CMs) between the LD and NMDA treatment (or between injury and development) were identified for each cell group by comparing chromVAR z-score with other groups for each condition. Motifs with an absolute log2 fold change (log2FC) greater than 0.25 and a adj-pvalue less than 0.05 were classified as CMs. Additionally, CMs were excluded if they were inconsistent with the list of CEGs, and also the highest correlation CMs were kept in the final list.

### Constructing gene regulatory networks using muti-omics datasets

In the paper, four GRNs were constructed as follows: 1) LD GRNs, which were constructed using only the samples from LD injury datasets. 2) NMDA GRNs, which were constructed using only the samples from NMDA injury datasets. 3) Injury GRNs, which were constructed employing the samples from both LD and NMDA injury datasets. 4) Development GRNs, which were constructed only using the samples from the development datasets.

### 1. Inferring activators and repressors by expression and motif activity

For each TF-motif pair in each datasets (LD,NMDA,Injury,Development), we calculate the Pearson correlation between TF expression and motif activities (chomVAR score) at single cell level with the “cor” function in R. Activator and repressor TF-motif pairs were identified if their correlation are large than 0.05 or less than −0.05.

### 2. Identifying Cis-regulatory elements

Firstly, the categorization of all peaks into three groups based on their genomic location relative to gene loci is performed: 1) Promoter (within 500bp of TSS), 2) Gene Body, and 3) Intergenic. Subsequently, peak-target pairs are generated using the following methods: 1) The target genes for Promoter and Gene Body peaks are determined by the genes they overlap with. 2) The target genes for Intergenic peaks encompass all genes located within 200kb of the peak’s location.

Next, PtoG correlations for each peak with its surrounding genes (200kb) are calculated using the “addPeak2GeneLinks” function in the ArchR package. The first 30 dimensions of the combined multi-omic space are utilized to generate cell groups using ArchR.

Finally, the retention of the peak-target pairs is based on meeting the following criteria for their PtoG correlations: abs(correlation) > 0.25 and FDR < 0.01.

### 3. Predicting TF Binding Sites

The TF-peak pairs were constructed by predicting TF binding sites inferred based on motif information and scATACseq footprint signals within the identified cis-regulatory elements.

Initially, Position Weight Matrices (PWMs) were extracted from the TRANSFAC2018 and CIS-BP databases. The binding regions were then identified by matching these motifs to the DNA sequences of the cis-regulatory elements using the motifmatchr package^79^ (’matchMotifs’, p.cutoff = 5e-05).

Subsequently, the scATACseq corrected footprint signals were separately calculated for the Light-damage Injury, NMDA injury, combined injury, and Development datasets. For the Light-damage and NMDA injury datasets, the insertion fragments from Rest MG, Activated MG, and MGPCs cells were combined. For the development datasets, the insertion fragments from MG, Early RPCs, and Late RPCs cells were merged. These merged fragments were converted to BAM format and processed through the TOBIAS pipeline^80^ to obtain bias-corrected Tn5 signals.

For each binding region of the motif, footprint scores were computed, including NC, NL, and NR. NC represented the average Tn5 signal at the center of the motif, while NL and NR indicated the average Tn5 signals in the left and right flanking regions (triple the size of the center) of the binding region, respectively.

Finally, the TF binding sites were retained based on the following criteria: NR+NL-2*NC > 0.1. Additionally, the binding regions for motifs whose corresponding TFs were not expressed in the MG cells were removed.

### 4. TF-target correlation

The TF-target relationship was calculated using the Stochastic Gradient Boosting Machine (SGBM) method, which was implemented through the arboreto package ^81^ in Python. The “grnboost2” function from the package took the expression matrix, gene list, and TF list as inputs, generating a table of TF-gene pairs with important scores. The TF-gene pairs were filtered based on their important scores, with pairs that had scores lower than the 95th quantile being removed. Additionally, the Pearson correlation between each TF-gene pair was computed according to the cell-by-gene expression matrix. If the correlation exceeded 0.03, the TF-gene pair was annotated as “positive” regulation; if the correlation was below −0.03, it was annotated as “negative” regulation. Any other TF-gene pairs were filtered out.

### 5. Construction of TF-peak-target links

The total GRNs for each condition (LD,NMDA,Injury,and Development) were constructed by integrating data from the previous steps. The following procedure was employed: The TF-peak pairs from step 3 and the peak-target pairs from step 2 were merged to form TF-peak-target triples. Subsequently, these TF-peak-target triples were filtered using the following criteria: 1) The triples were retained only if TF activity are in the same direction with TF-gene correlation. (Activtor with positive TF-gene correlation, Repressor with negative TF-gene correlation) 2) The triples were retained only if the TF’s expression levels are enriched in MG cell groups (MG, ActMG, MGPCs). 3) Any duplicate triples were eliminated, and we retained the highest footprint score for each TF-peak-target pair.

### 6. Identification of enriched gene regulatory sub-networks

The enriched sub-GRNs were extracted from the total GRNs generated in step 5, based on the logFC change of TFs, peaks, and target genes (as shown in Figure 4E and Figure 6E). For instance, to obtain enriched LD sub-GRNs, we applied the following criteria to filter triple pairs (TF-Peak-Target) from the total LD GRNs: 1) The expression levels of TFs should be higher (or lower, if TF-motif pairs are repressors) in LD (compared to NMDA) in at least one of the MG, ActMG, and MGPCs cell groups comparisons. 2) The accessibility of peaks should not be lower in LD (compared to NMDA) in any of the MG, ActMG, and MGPCs cell groups. 3) The expression levels of target genes should be higher in LD (compared to NMDA) in at least one of the MG, ActMG, and MGPCs cell groups comparison. The same methods were employed to extract sub-GRNs from NMDA GRNs, Injury GRNs, and Development GRNs.

### 7. Identication of key activator TFs

To identify the key activators (TFs) in the GRNs, we initially reduce the triple pairs (TF-peak-target) into double pairs (TF-target) for each GRNs. For each TF in the network, we first calculate coverage score to see how many DEGs are regulated by that TF. For example, to identify key activators in LD enriched sub-network, for each TF, we first count the number of overlap genes between its targets with LD enriched DEG clusters (cluster 1,2,and 3 in Figure 4G). For each TF and each DEG cluster, the coverage were calculated as: N_overlap_ / N_cluster_, where N_overlap_ denote as the number of overlapped genes, and N_cluster_ denote as the total number of DEGs in that cluster. Next, to test whether the TFs are significantly regulates the DEG clusters, we used hypergeometric test with “phyper” function in R for each TF and DEG cluster.

In the hypergeometric test, the “population” is defined as the TF target genes from total GRNs. The “sample” is defined as the TF target genes from LD enriched sub-GRNs and the “successes” is defined as the genes from the DEGs cluster. Finally, all the TFs with p-value < 0.001 and coverage > 0.01 were identified as key activators.

### ChromVAR analysis

A ChromVAR ^82^ analysis was performed to determine the global transcription factor (TF) activity in each cell. The raw cell-by-peak matrix was initially inputted into ChromVAR, and the “addGCBias” function was used to correct for GC bias. The DarRer11 reference genome was employed for this correction.

Subsequently, a TF z-score matrix was created by combining motifs from the TransFac2018 and CIS-BP databases. This was achieved using the “matchmotifs” and “computeDeviations” functions. The z-score of each cell was then utilized to generate a heatmap and visualization. These outputs were overlaid onto the previously calculated UMAP coordinates.

### GO analysis

To identify significantly enriched Gene Ontology (GO) terms (specifically Biological Process) and KEGG terms among the Differentially Expressed Genes (DEGs) between biological conditions, the gene set enrichment analysis was performed using the “enrichGO” and “enrichKEGG” functions from the “clusterProfiler” R package ^83^.

## Supplemental Tables/Datasets

**Table ST1:** Total cell counts for LD and NMDA.

**Tabel ST2**: snRNA/ATAC-Seq and scRNA-Seq data for LD and NMDA samples.

**Table ST3:** Genes, peaks, and motifs selectively enriched in LD vs. NMDA samples (snRNA/ATAC-Seq+scRNA-Seq).

**Table ST4:** Regulatory relationships among TFs selectively active in LD vs. NMDA samples.

**Table ST5:** snRNA/ATAC-Seq and reanalyzed scRNA-Seq data for developing retinal samples.

**Table ST6:** Genes, peaks, and motifs selectively enriched in injury vs. development samples (snRNA/ATAC-Seq+scRNA-Seq).

**Table ST7:** Regulatory relationships among TFs selectively active in development vs. injury samples.

## SUPPLEMENTAL FIGURES

**Supplemental Figure 1:**
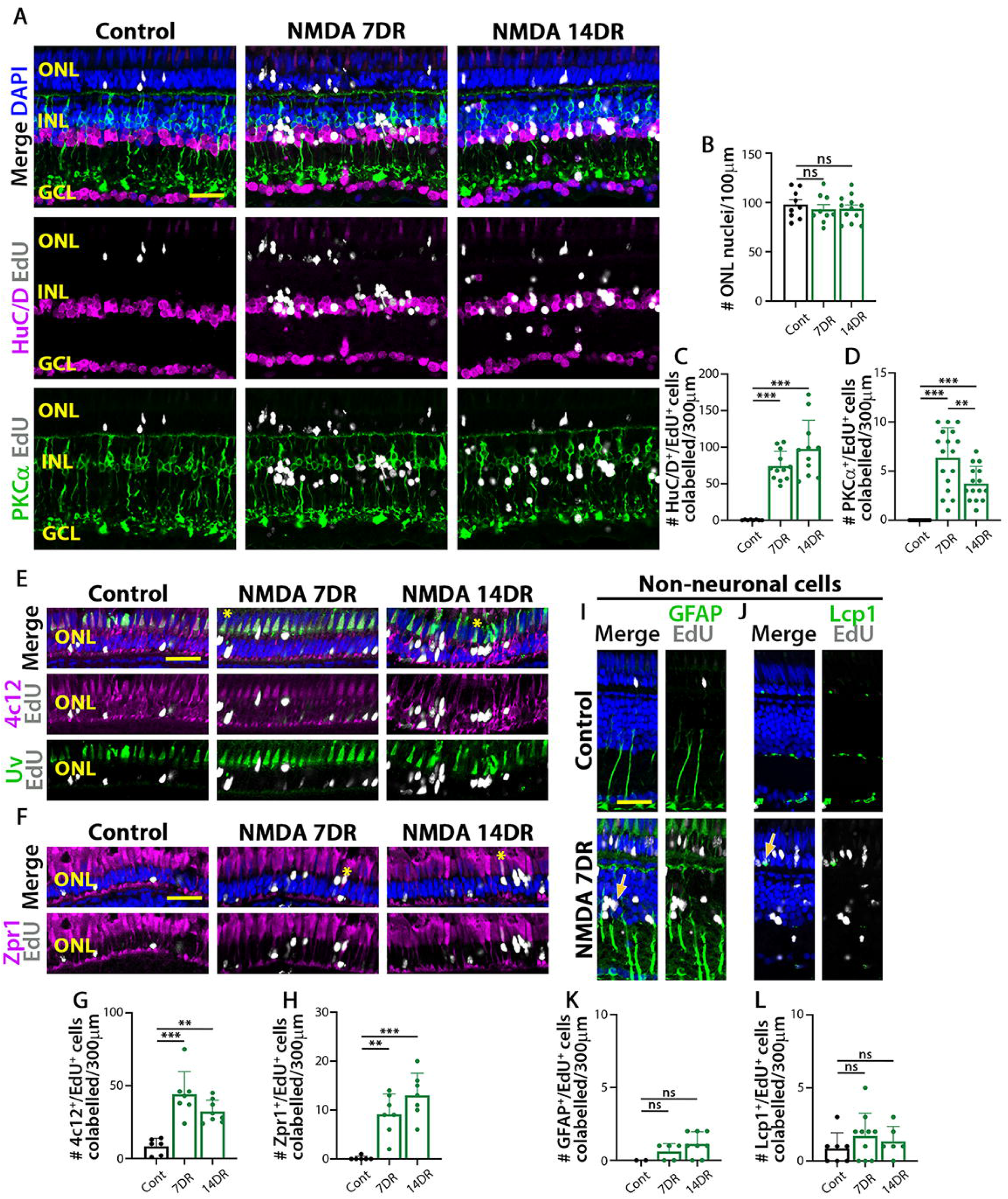
Regeneration of retinal neurons following NMDA damage. (A) EdU-labeling in retinas prior to NMDA damage (Control) and 7 and 14 days following 96 hours of NMDA damage. Retinas were immunostained for EdU, HuC/D, and PKCa and counterstained with DAPI. (B) Quantification of the number of DAPI-labeled nuclei in the INL. (C) Quantification of the number of cells colabeled for EdU and HuC/D. (D) Quantification of the number of cells colabeled for EdU and PKCa. (E) EdU-labeling in retinas prior to NMDA damage (Control) and 7 and 14 days following 96 hours of NMDA damage. Retinas were immunostained for EdU, 4c12, and UV opsin and counterstained with DAPI. (F) EdU-labeling in retinas at 7 and 14 days following 96 hours of NMDA damage. Retinas were immunostained for EdU and Zpr1 and counterstained with DAPI. (G) Quantification of the number of cells colabeled for EdU and 4c12. (H) Quantification of the number of cells colabeled for EdU and Zpr1. (I) EdU-labeling in retinas prior to NMDA damage (Control) and 7 days following 96 hours of NMDA damage. Retinas were immunostained for EdU and GFAP and counterstained with DAPI. (J) EdU-labeling in retinas prior to NMDA damage (Control) and 7 days following 96 hours of NMDA damage. Retinas were immunostained for EdU and Lcp1 and counterstained with DAPI. (K) Quantification of the number of cells colabeled for EdU and GFAP. (L) Quantification of the number of cells colabeled for EdU and Lcp1. Scale bars in A, E, F, I, and J are 20μm.

**Supplemental Figure 2:**
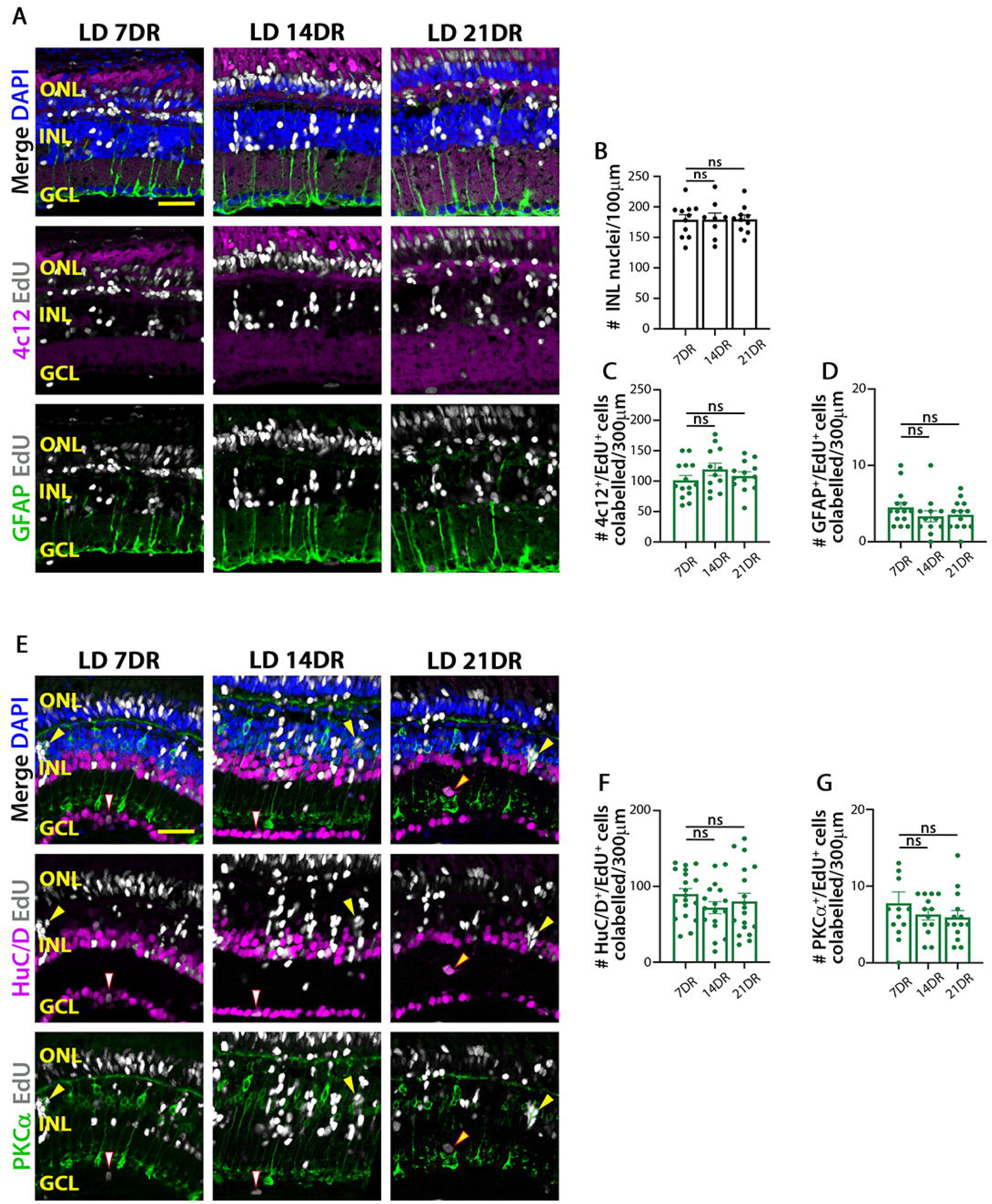
Regeneration of retinal neurons following constant light damage. (A) EdU-labeling in retinas at 7, 14, and 21 days following 96 hours of constant light. Retinas were immunostained for EdU, 4c12, and GFAP and counterstained with DAPI. (B) Quantification of the number of DAPI-labeled nuclei in the INL. (C) Quantification of the number of cells colabeled for EdU and 4c12. (D) Quantification of the number of cells colabeled for EdU and GFAP. (E) EdU-labeling in retinas at 7, 14, and 21 days following 96 hours of constant light. Retinas were immunostained for EdU, HuC/D, and PKCa and counterstained with DAPI. (F) Quantification of the number of cells colabeled for EdU and HuC/D. (G) Quantification of the number of cells colabeled for EdU and PKCa. Scale bars in A and E are 20μm.

**Supplemental Figure 3:**
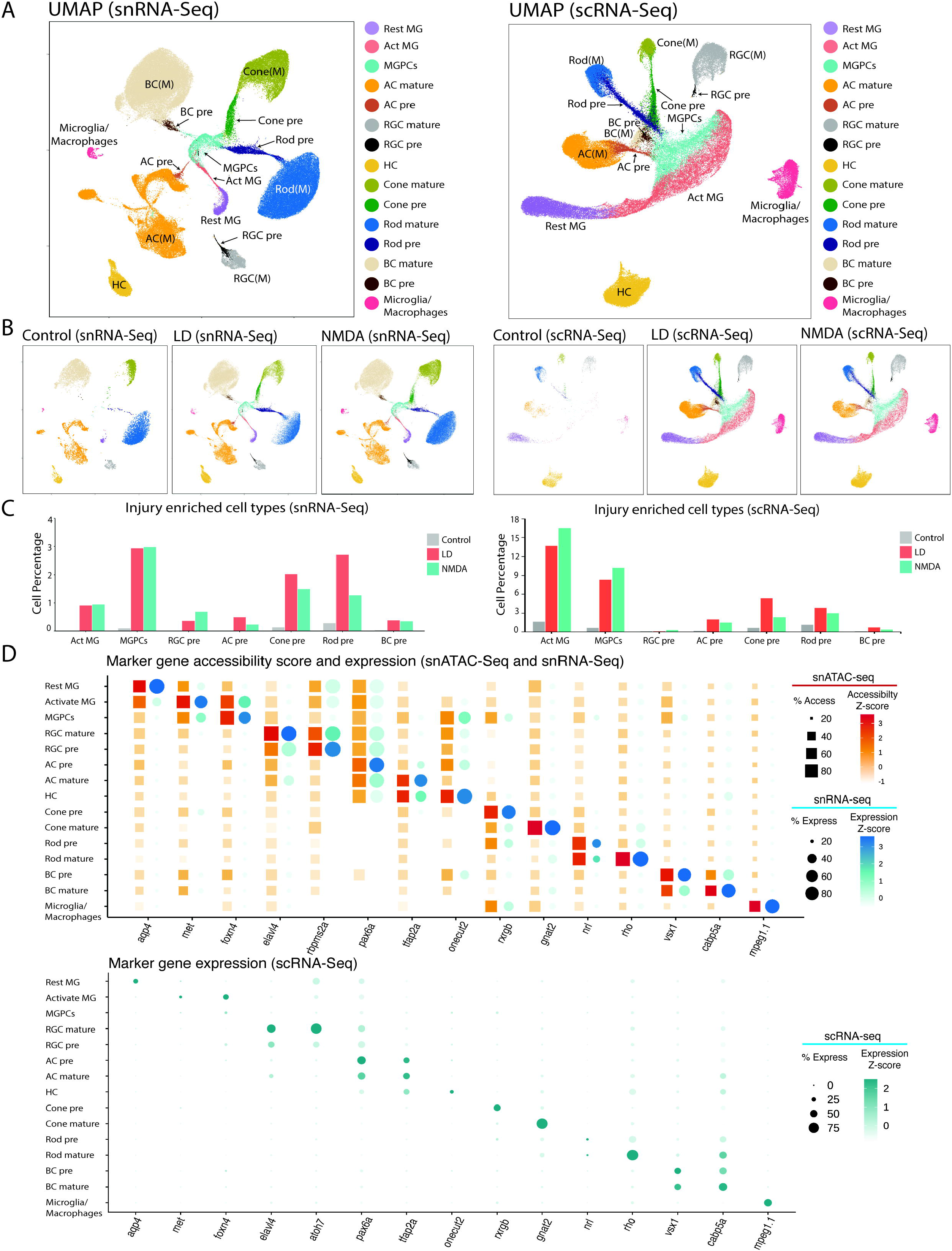
Overview of single-cell sequencing datasets from LD and NMDA injury models. (A) Combined UMAP plots showing the cells in LD and NMDA datasets (left: snRNA-seq, right: scRNA-seq). Each point represents an individual cell and is colored by its corresponding cell type. (B) UMAP plots (left:snRNA-seq, right:scRNA-seq) showing the cells separated by each condition for both snRNA-seq and scRNA-seq datasets. (C) The bar plot (left:snRNA-seq, right:scRNA-seq) indicates Act MG, MGPCs and newly-borned neuron precursors are both enriched in the two injury models compared to control dataset. The y axis indicated the cell ratio at each time point. Bars are colored by conditions. (D) Examples of mRNA levels and chromatin accessibility (up panel:snRNA-seq and snATAC-seq, down panel:scRNA-seq) for selected cell-type-specific genes.

**Supplemental Figure 4:**
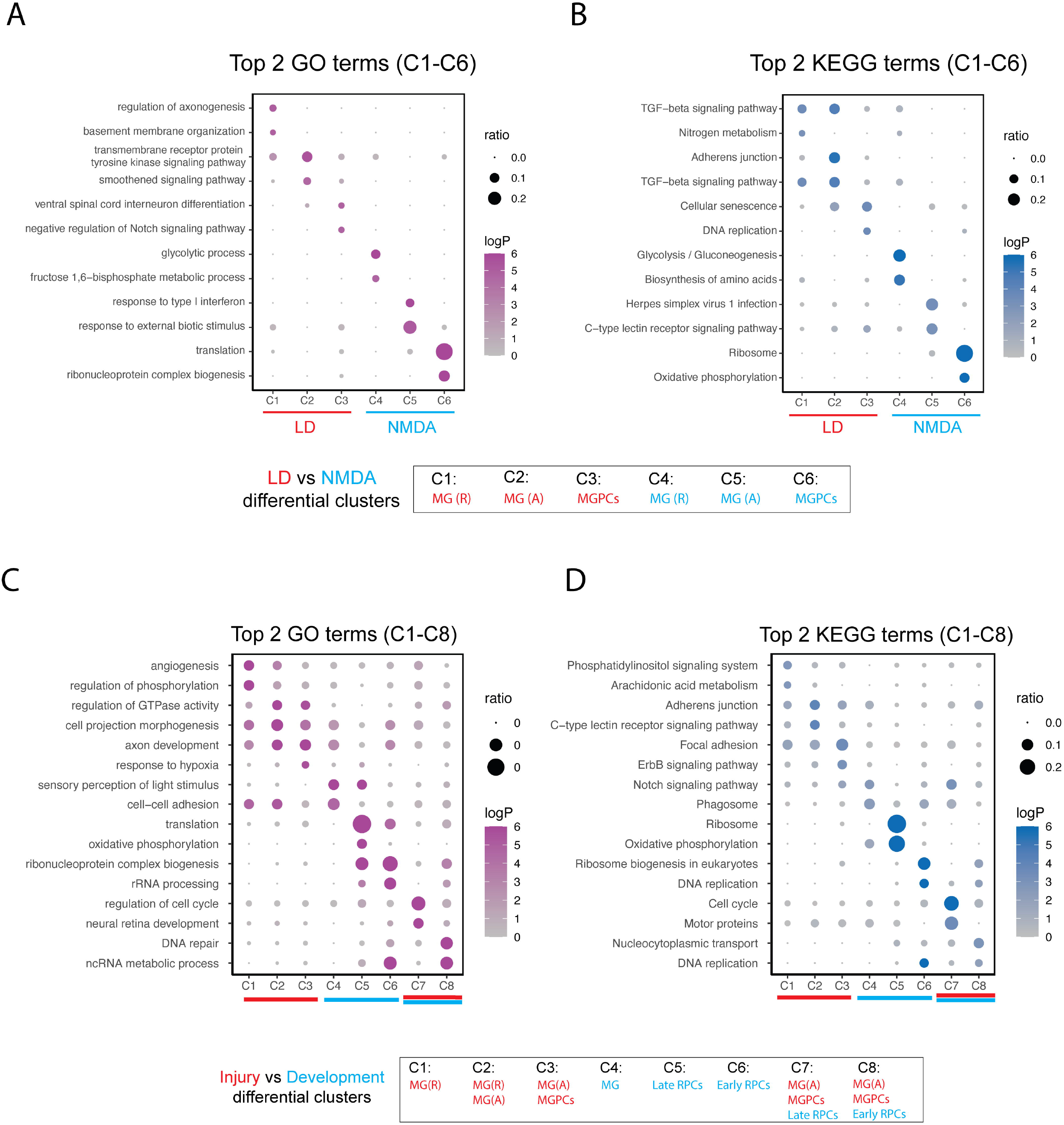
Enriched GO and KEGG terms in LD and NMDA, and injury and development. (A,B) Dot plots showing the most highly enriched 2 GO and KEGG terms enriched for differential gene clusters between LD and NMDA models. The x-axis indicates the clusters and the y-axis indicates the GO or KEGG terms, the size of the dot represents the gene ratio and the color represents the negative log-transformed p-values. (C,D) Dot plots showing the most highly enriched 2 GO and KEGG terms enriched for differential gene clusters between injury and development models. The x-axis indicates the clusters and the y-axis indicates the GO or KEGG terms, the size of the dot represents the gene ratio and the color represents the negative log-transformed p-values.

**Supplemental Figure 5:**
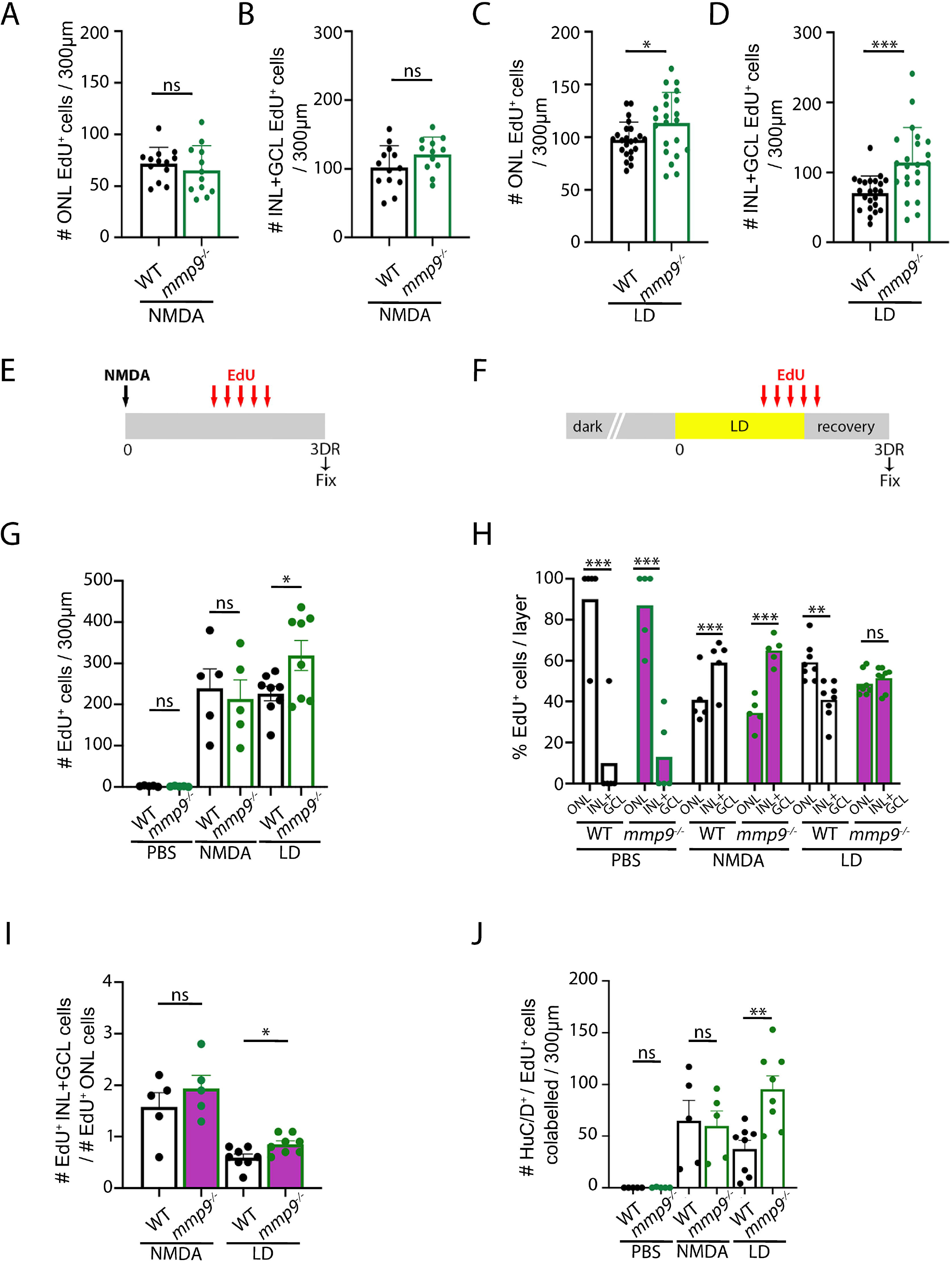
Mmp9 is required for regulating the regeneration of HuC/D-labeled amacrine and ganglion cells. (A) Quantification of the number of EdU-labeled ONL cells in NMDA-injected wild-type and *mmp9* mutant retinas at 7 DR. (B) Quantification of the number of EdU-labeled INL+GCL cells in NMDA-injected wild-type and *mmp9* mutant retinas at 7 DR. (C) Quantification of the number of EdU-labeled ONL cells in light-damaged wild-type and *mmp9* mutant retinas at 7 DR. (D) Quantification of the number of EdU-labeled INL+GCL cells in light-damaged wild-type and *mmp9* mutant retinas at 7 DR. (E) Schematic of NMDA-induced damage experiment. (F) Schematic of light-induced damage experiment. (G) Quantification of the number of EdU-labeled cells in all three retinal layers in wild-type and *mmp9* mutants following either PBS injection, NMDA damage, or light damage at 3 DR. (H) The percentage of EdU-positive cells in the ONL versus the INL+GCL is plotted for wild-type and *mmp9* mutant retinas after PBS injection, NMDA damage, and light damage at 3 DR. (I) The ratio of EdU-positive INL+GCL cells to EdU-positive ONL cells is plotted for wild-type and *mmp9* mutant fish following either NMDA damage or light damage at 3 DR. (J) Quantification of the number of cells colabeled for EdU and HuC/D in PBS-injected, NMDA-damaged, and light-damaged retinas at 3 DR.

**Supplemental Figure 6:**
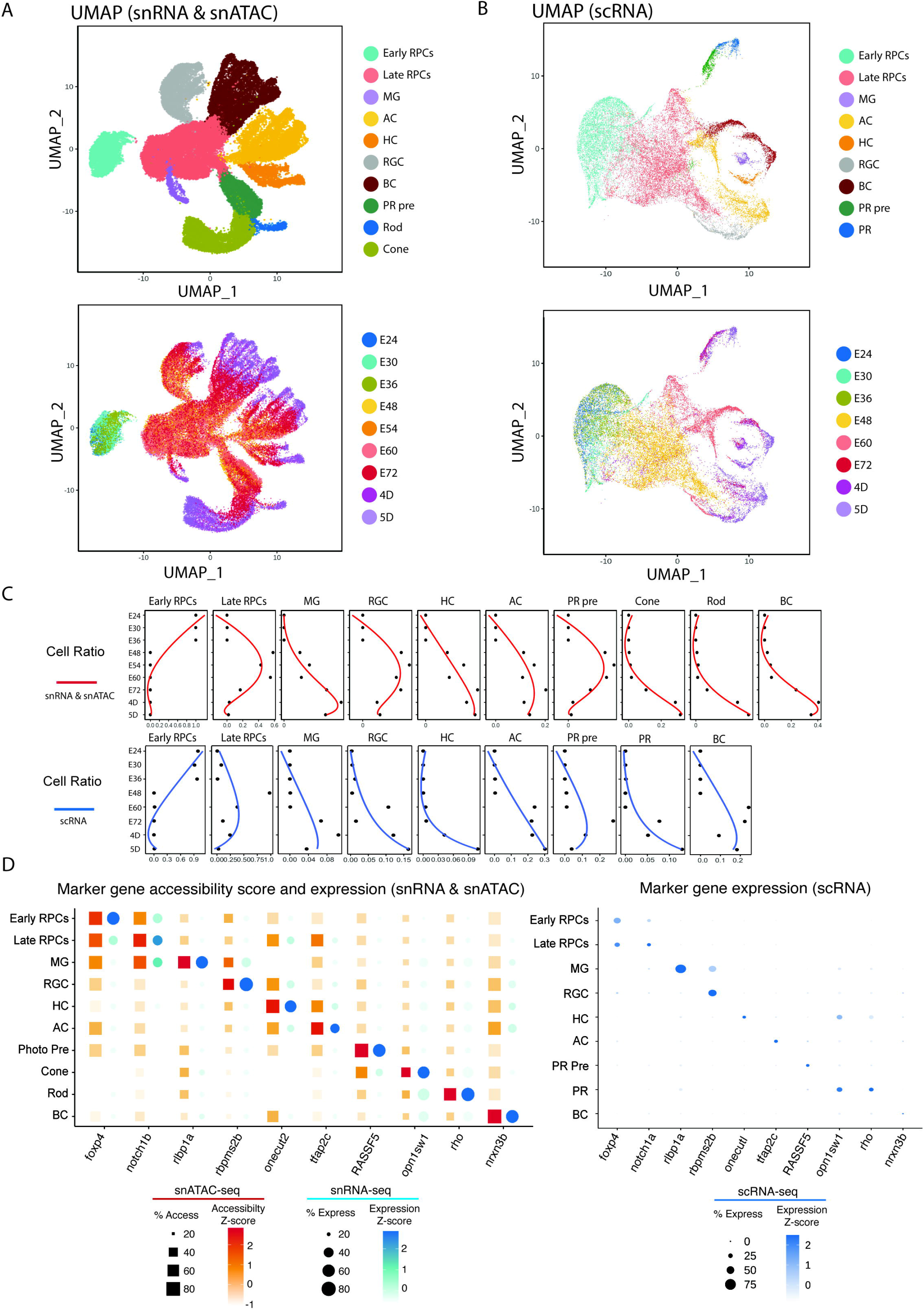
Overview of the single-cell sequencing datasets of the development model. (A) UMAP plots showing cells present in snRNA-seq datasets from developing retina. Each point is colored by cell type (up panel) and time point (down panel). (B) UMAP plots showing cells present in scRNA-seq datasets from developing retina. Each point is colored by cell type (up panel) and time point (down panel). (C) The line plot (up panel:snRNA-seq, down panel:scRNA-seq) showing the changing of cell ratios during the development. X-axis indicates the cell ratio and y-axis indicates each time point. (D) Examples of mRNA levels and chromatin accessibility (left:snRNA-seq and snATACseq, right:scRNA-seq) for selected cell-type-specific genes.

**Supplemental Figure 7:**
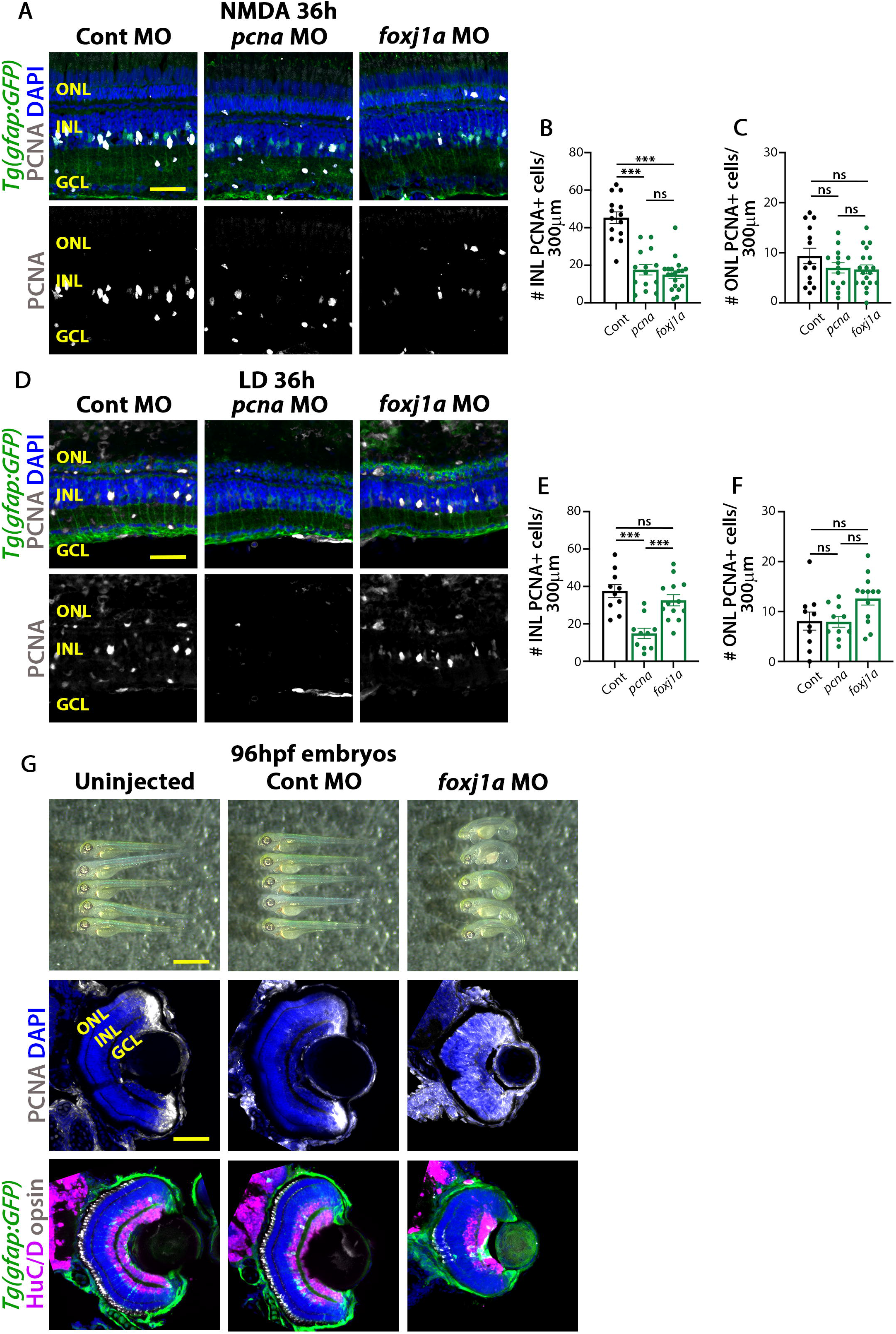
Foxj1a is required for MGPC proliferation. (A) Tg(*gfap:GFP*) retinas electroporated with either Standard Control morpholino (Cont MO), *pcna* MO, or *foxj1a* MO were isolated 36 hours after NMDA injection and immunostained for PCNA, GFP, and counterstained with DAPI. (B) Quantification of the number of PCNA-labeled cells in the INL. (C) Quantification of the number of PCNA-labeled cells in the ONL. (D) Tg(*gfap:GFP*) retinas electroporated with either Cont MO, *pcna* MO, or *foxj1a* MO were isolated 36 hours after starting constant light and immunostained for PCNA, GFP, and DAPI. (E) Quantification of the number of PCNA-labeled cells in the INL. (F) Quantification of the number of PCNA-labeled cells in the ONL. (G) Uninjected and Standard Control morphants, as well as *foxj1a* morphants, were examined for gross morphological phenotypes at 96 hpf. Uninjected, Standard Control, and *foxj1a* morphant sections retinas were stained for PCNA and DAPI. In addition, *Tg(gfap:GFP)* uninjected, Standard Control, and *foxj1a* morphant retinas at 96 hpf were immunostained for HuC/D, green opsin (double cone photoreceptors), GFP, and counterstained with DAPI. Scale bars in A and D are 20μm, in G (top) is 1 millimeter, and in G (middle) is 50μm.

